# Zebrafish *dazl* regulates cystogenesis upstream of the meiotic transition and germline stem cell specification and independent of meiotic checkpoints

**DOI:** 10.1101/2019.12.23.887687

**Authors:** Sylvain Bertho, Mara Clapp, Torsten U. Banisch, Jan Bandemer, Erez Raz, Florence L. Marlow

## Abstract

Fertility and gamete reserves are maintained by asymmetric divisions of the germline stem cells to produce new stem cells or daughters that differentiate as gametes. Before entering meiosis, differentiating germ cells (GCs) of sexual animals typically undergo cystogenesis. This evolutionary conserved process involves synchronous and incomplete mitotic divisions of a germ cell daughter (cystoblast) to generate sister cells connected by stable intercellular bridges that facilitate exchange of materials to support a large synchronous population of gamete progenitors. Here we investigate cystogenesis in zebrafish and identified Deleted in azoospermia (Dazl), a conserved vertebrate RNA binding protein as a regulator of this process. Analysis of *dazl* mutants revealed an essential role for Dazl in regulating incomplete cytokinesis and germline cyst formation before the meiotic transition. Accordingly, *dazl* mutant GCs form defective ring canals, and ultimately remain as individual cells that fail to differentiate as meiocytes. In addition to promoting cystoblast divisions and meiotic entry, *dazl* function is required upstream of germline stem cell establishment and fertility.

**Summary Statement:** We show that zebrafish *dazl* is required for incomplete cytokinesis to generate germline cysts during cystogenesis, acts upstream of germline stem cell establishment, and is required for meiosis, and fertility.

## Introduction

In many organisms the germline is among the first cell types to be set aside during early development (Ginsburg, 1994; Illmensee and Mahowald, 1976; Illmensee et al., 1976; Wolf et al., 1983). Early germ cells, called primordial germ cells (PGCs), can be specified by maternal factors or by inductive signals (Farrell et al., 2018; Lawson and Hage, 1994; Marlow, 2015; Nieuwkoop and Sutasurya, 1979). Once specified, PGCs ignore somatic differentiation programs and migrate to the site where the gonad anlage forms (Braat et al., 1999a; Gross-Thebing et al., 2017; Nieuwkoop and Sutasurya, 1979; Strome and Updike, 2015). There the PGCs proliferate and eventually enter meiosis and differentiate to produce the gametes, sperm in males and oocytes in females. Although the earliest stages of PGC development in zebrafish have been extensively studied (Barton et al., 2016; Marlow, 2015; Paksa and Raz, 2015; Raz, 2003), the cellular events and molecular mechanisms involved in the transition from PGC to germline stem cell (GSC) and the earliest phases of gametogenesis are less well understood.

### Germline cyst formation

A common and evolutionarily conserved feature of GCs is the asymmetric division of GSCs to produce a stem cell and a premeiotic daughter. These differentiating divisions have been classified as type-I or type-II divisions (Saito et al., 2007; Saito and Tanaka, 2009). Cells resulting from type-I divisions do not divide further but directly differentiate as meiotic cells and are observed in juvenile and adult teleost fish (medaka and in zebrafish)(Marlow and Mullins, 2008; Saito et al., 2007). Type-II divisions generate cystoblast cells that divide mitotically with incomplete cytokinesis to generate interconnected sisters in *Drosophila* (Cox and Spradling, 2003), *Xenopus* (Kloc et al., 2004) and medaka (Saito et al., 2007). The interconnections generated from incomplete cytokinesis form intercellular bridges (or ring canals) (Brown and King, 1964; de Cuevas et al., 1997; Fawcett et al., 1959; Greenbaum et al., 2011; Haglund et al., 2011; Koch and King, 1966, 1969; Koch et al., 1967; Lei and Spradling, 2016; Lin and Spradling, 1993; Mahowald, 1971; Marlow and Mullins, 2008; Pepling and Spradling, 1998; Robinson and Cooley, 1996; Spradling, 1993). In *Drosophila* ring canal formation involves regulation of the actin cytoskeleton, which localizes to the cleavage furrows and maintains midbody structures (Greenbaum et al., 2011). Arrest of the cytokinetic furrow and maintenance of the midbody involves a complex regulatory network that stabilizes the actomyosin meshwork to arrest abscission and maintain the contractile ring, forming ring canals instead of dividing (Greenbaum et al., 2011; Haglund et al., 2011; Hime et al., 1996; Robinson and Cooley, 1996). In mouse, interaction between the inactive serine-threonine kinase TEX14 and CEP55 regulates intercellular bridge stability in part by blocking abscission factors (Alix, Escrt complex) (Greenbaum et al., 2011; Greenbaum et al., 2006; Kim et al., 2015; Morita et al., 2007). Intercellular bridge and germline cyst formation is a conserved feature of germ cell biology and is crucial for fertility (e.g. (Greenbaum et al., 2006)). Intercellular bridges of germ cells are thought to fulfill several functions, including facilitating intercellular communication to produce numerous synchronous cells (de Cuevas et al., 1997; Lei and Spradling, 2013; Pepling and Spradling, 1998), coordinating critical stages such as meiotic entry (Robinson and Cooley, 1996; Stanley et al., 1972), maintenance of “functional ploidy” and gamete equivalence after meiotic divisions, and a mechanism for sensing and selection against abnormalities that would be detrimental to subsequent generations (Braun et al., 1989; LeGrand, 2001).

### Dazl and fertility

The RNA binding protein (Rbp) Deleted in azoospermia-like (Dazl) is a member of the Deleted in azoospermia (DAZ) family composed of Daz, Daz-like (Dazl) and Boule (Fu et al., 2015). DAZ family members are exclusively expressed in the germline and contribute to various aspects of GC development in invertebrates and vertebrates (Fu et al., 2015). Although *dazl* is required for GC development, loss of *daz* family members disrupts distinct aspects of GC development in different species (Alphey et al., 1992; Courtot et al., 1992; Eberhart et al., 1996; Fu et al., 2015; Fukuda et al., 2018; Gill et al., 2011; Iyer et al., 2016; Karashima et al., 2000; Maines and Wasserman, 1999; Ruggiu et al., 1997; Saunders et al., 2003; Schrans-Stassen et al., 2001). In early embryos, antisense oligodeoxynucleotide depletion of *Xenopus* Dazl disrupts PGC migration and causes PGC deficiency (Houston and King, 2000). Recently, Dazl has been shown to regulate RNAs involved in the meiotic program (Haston et al., 2009b; Kim et al., 2012; Li et al., 2019; Medrano et al., 2012; Zagore et al., 2018) and very recently has been shown to regulate commitment to germline fate (Nicholls et al., 2019). These studies identify diverse and opposing activities for Dazl including regulation of RNA stability, translational activation and translational repression of its target RNAs (Li et al., 2019; Zagore et al., 2018). For example, in mouse testis Dazl is implicated in translational repression of pluripotency genes (Chen et al., 2014). In zebrafish PGCs, overexpression studies indicate Dazl antagonizes miR-430 and promotes polyadenylation of maternally-provided germline RNAs (Maegawa et al., 2002; Takeda et al., 2009); however, Dazl’s essential role in germline development in zebrafish remains unknown.

Our study examines the earliest stages of gonadogenesis and provides evidence that Dazl is required for germline cyst formation and plays critical roles in GC amplification upstream of meiotic entry and establishment of GSCs, and thus is essential for fertility. Here we describe zebrafish cystogenesis, a process during which PGCs transit from individual cells to GC clusters that undergo morphological changes characterized by complex cytoplasmic and nuclear rearrangements to form germline cysts. We show that GC numbers increase concomitantly with cyst formation. Following F-Actin distribution revealed that premeiotic cyst cells are interconnected by intercellular bridges or ring canals and contain aggregated or branched actin rich structures reminiscent of spectrosomes and fusomes (Cooley and Theurkauf, 1994; Deng and Lin, 1997; Lighthouse et al., 2008; Lin et al., 1994; Robinson and Cooley, 1996, 1997; Tilney et al., 1996; Xue and Cooley, 1993; Yue and Spradling, 1992). Analysis of zinc finger nucleases and CRISPR-induced *dazl* mutant alleles revealed that zygotically transcribed *dazl* is essential for incomplete cytokinesis and germline cyst formation before meiosis begins. In contrast to wild-type GC, *dazl* mutant GCs form defective ring canals, and ultimately remain as individual cells that fail to differentiate as meiocytes. In addition to promoting cystoblast divisions and meiotic entry, Dazl acts upstream of GSC establishment, and consequently *dazl* mutant fish develop exclusively as sterile males. Zygotic *dazl* is however dispensable for GC specification, germ granule formation, GC migration, and PGC divisions.

In addition, we show that the apoptotic pathway mediated by the *cell cycle checkpoint kinase*, *chk2,* is dispensable for early development, sex determination, gonadogenesis, and fertility in zebrafish. Although germline survival was prolonged in *dazl* mutants devoid of *chk2,* loss of *dazl^-/-^* GCs was not suppressed by *tp53* or *chk2* mutations, indicating that germline loss occurs via an independent mechanism. Our findings support a novel requirement for *dazl* in GCs for cystogenesis upstream of GSC establishment prior to the meiotic transition and provide evidence that type-II or amplifying divisions and cyst formation are critical for the mitotic to meiotic transition and fertility in zebrafish.

## Materials and methods

### Fish strains

*dazl^ae57^* and *dazl^ae34^* mutant fish strains were generated using Crispr-Cas9 mutagenesis as in (Gagnon et al., 2014). *dazl^Δ7^* was generated using zinc fingers nucleases (Ekker, 2008; Foley et al., 2009b) as detailed below. Complementation tests were performed by intercrossing carriers of each *dazl* mutant allele. To generate double mutants, *dazl^ae57^* was crossed with *tp53^M214K^* or the *chk2^sa20350^* allele, generated in The Sanger Institute’s Zebrafish Mutation Project (Kettleborough et al., 2013) and obtained through ZIRC. Genomic DNA and cDNA from *chk2 ^sa20350^* mutant tissues were sequenced to verify the genomic mutation and determine the transcript produced from the mutant allele (primers in Supplemental Table 1). To visualize the GCs, mutations were crossed into the *ziwi:GFP* GC reporter line (Leu and Draper, 2010). All procedures and experimental protocols were performed in accordance with NIH guidelines and were approved by the Einstein (protocol #20140502) and Icahn School of Medicine at Mount Sinai Institutional (ISMMS) Animal Care and Use Committees (IACUC #2017-0114).

### dazl genotyping

Genomic DNA (gDNA) was obtained either from dissected adult trunk, fin clip, or whole larvae. Samples were lysed in an alkaline lysis buffer (25 mM NaOH, 0.2 mM EDTA, pH12), heated at 95°C for 20 mins. Then, cooled to 4°C before neutralizing buffer was added (20 mM Tris-HCl, 0.1 mM EDTA, pH 8.1) (Truett et al., 2000). gDNA from *dazl^ae57^* or *dazl^ae34^* was PCR-amplified for 40 cycles with an annealing temperature of 59°C, followed by an Eco147I restriction enzyme digestion for 1 hour (fast digest, Eco147I, Thermo scientific). Undigested and digested products were resolved in a 3% Metaphor 1:1 (lonza)/agarose gel (Invitrogen) gel. Eco147I cuts the WT fragment. A PCR amplified fragment annealed at 57°C flanking the *dazl^Δ7^* region was digested with AflIII enzyme then resolved on a 3% Metaphor/agarose gel. AflIII cuts the mutant fragment.

*tp53^M214K^* was identified as previously described (Berghmans et al., 2005). High resolution melt analysis (HRMA) assays were also developed for *dazl^ae57^*, *dazl^ae34^*, *tp53^M214K^*, *chk2^sa20350^,* and the *dazl^delta7^* (described below) alleles (primers in (Supplemental Table1)).

### Mutagenesis

*dazl^ae34^* and *dazl^ae57^* allele were generated by CRISPR-Cas9 mediated mutagenesis based on (Gagnon et al., 2014). *dazl* gRNAs targeting exon 6 were designed using the CHOPCHOP webtool (Montague et al., 2014) (Supplemental Table1). Briefly, the gene specific target and the constant oligonucleotides were annealed, and the fragment was filled in using T4 DNA polymerase. Next, the fragment was transcribed and cleaned up to yield sgRNA using the MEGAscript SP6 kit (Life technology, Ambion). 1 nl of 12.5 ng/µl of gRNA and 1 nl of Cas9 protein (300 ng/ul) (Jinek et al., 2012) along with phenol red (Sigma Aldrich) were co-injected at one-cell stage. At 24 hpf, uninjected and injected embryos (n=8 each) were assayed by PCR amplification (primers in Supplemental table 1) followed by T7 endonuclease digest (Hwang et al., 2013), and those with new banding patterns were sequenced to confirm mutagenesis. Injected embryos were raised to adulthood, and individuals carrying mutations were identified by extracting gDNA from their progeny and fin tissue and assaying as above. Smaller bands compared to the WT allele, indicative of *de novo* mutations, were extracted from the gel, cloned into a PCR4 TOPO vector (Sigma), and sequenced in both directions to determine the mutated sequence. Fish harboring *dazl^ae34^* or *dazl^ae57^* mutations were outcrossed to AB fish. All mutations were verified by sequencing both genomic DNA and cDNA from mutant animals. Total RNA was extracted from pooled embryos from heterozygote intercrosses or AB strain WT (n=20-30) using Trizol (Life Technologies, 15596). cDNA was prepared with SuperScript III/IV Reverse Transcription Kit (Life Technologies, 18080-051). RT-PCR was performed to amplify the *dazl* coding region using Easy-A High Fidelity Taq polymerase (600400, Agilent) (primers in Supplemental Table 1). The PCR fragments were TOPO cloned into pCR8/GW/TOPO (K250020, Invitrogen) and sequenced (Macrogen). Sequences were analyzed using Sequencher or MacVector software.

The *dazl^Δ7^* allele, a 7nt deletion resulting in a frame shift and subsequent premature stop codon at amino acid 54, was generated using Zinc finger nucleases (ZFNs) (Foley et al., 2009a). The *dazl* genomic region was PCR amplified and sequenced. ZFNs targeting the region in exon two just upstream of the RRM were purchased from Sigma and injected (500pg) at one-cell-stage into wild-type embryos.

*dazl* ZFN set: left and right ZFN arrays in bold, spacer is underlined

GACGCCCAACAC**<u>ACTGT</u>**TCGTCGGCGGTATTG

Genomic lesions induced by ZFNs were subsequently identified by PCR amplification and RFLP analysis to identify founders carrying mutations in the germline. Recovered alleles were confirmed by sequencing.

PCR primers to identify lesions were:

5’ GTTCAGTTACCCGTGTGCCTGATAT 3’

3’ ATTAAACATTATAGTCCAATTAAACTAATCTGCATCA 5’

### Dissections

Fish were anesthetized with a lethal dose of tricaine (MS-22) and dissected. Fish were positioned laterally, and antero-posterior body length was measured prior to dissection. Images were acquired using an Olympus SZ61 dissecting microscope equipped with a high-resolution digital camera (model S97809, Olympus America) and Picture Frame 2.0 software.

### Immunostaining

For whole-mount immunofluorescence stained embryos or ovaries, tissues were fixed in 3.7% paraformaldehyde overnight at 4°C. The next day samples were washed in PBS then dehydrated in MeOH and stored at −20°C. Chicken anti-Vasa antibody (Blokhina et al., 2019) was used at a 1:5000 dilution. Alexafluor488, Alexafluor568, CY3, C5 (Molecular Probes) secondary antibodies were diluted 1:500. For F-actin labeling, samples were fixed overnight at 4°C in 3.7% paraformaldehyde in Actin Stabilization Buffer (Becker and Hart, 1999). Then, permeabilized in 2% Triton X-100. Following the primary antibody solution, Alexa fluor 568 phalloidin was added. Samples were mounted in vectashield with DAPI and images were acquired using a Zeiss Axio Observer inverted microscope equipped with ApotomeII and a CCD camera, a Zeiss Zoom dissecting scope equipped with ApotomeII, or a Leica SP5 DMI at the Microscopy CoRE at the IMSSM. Image processing was performed in Zenpro (Zeiss), Leica Application Suite (LAS), ImageJ/FIJI, Adobe Photoshop and Adobe Illustrator, Imaris (Oxford instruments).

### Germ cell, cyst and oocyte-like cell quantification

Vasa protein was used as a marker to identify and count individual GCs. Z-series stacks of gonads at each stage were obtained using a Zeiss Axio Observer inverted microscope equipped with ApotomeII and a CCD camera, or Zeiss Zoom dissecting scope equipped with ApotomeII, or a Leica SP5 DMI at the Microscopy CoRE at the ISMMS. Cells were manually counted by analyzing each slice within each Z-stack for Vasa positivity and nuclear morphology (DAPI) defining each category during the cystogenesis process and the number of cysts per cell. Quantification of spectrosomes was performed by counting each actin-rich structure through the Z-stack. Cell area and volume was measured after a manual segmentation of the cell in each plane through the Z-stack. All the above experiments were performed using ZEN pro (Zeiss), Leica Application Suite (LAS) or ImageJ/FIJI, Imaris (Oxford instruments).

### Statistical analysis

Statistical differences were assessed using Prism software and paired Student’s t-test. Significant differences are indicated in figures by asterisks if p<0.001 (***), p<0.05 (**), p<0.01 (*) or n.s. if not significant.

## Results

### Formation of germline cysts

Following their arrival at the gonad, PGCs proliferate before initiating meiosis to form a bipotential gonad comprised of oocyte-like cells (OLCs) (Leu and Draper, 2010; Tong et al., 2010; Tzung et al., 2015; Wang et al., 2007). Timing of entry into meiosis and OLC abundance are thought to contribute to sexual fate determination, such that more GCs promote female fate (Ye et al., 2019). Conversely, low numbers of germ cells and OLCs drive male development (Dai et al., 2015; Orban et al., 2009; Tzung et al., 2015). Despite these correlations highlighting the importance of proper development of OLCs, little is known about the actual mechanisms and cell biology regulating GC numbers and meiotic entry in zebrafish. Type-II divisions of GSCs have been observed in juvenile and adult teleosts (medaka and in zebrafish) and are thought to be amplifying divisions (Beer and Draper, 2013; Marlow and Mullins, 2008; Saito et al., 2007). In zebrafish this process likely occurs between 5-14 days when PGCs proliferate before initiating meiosis (Leerberg et al., 2017; Leu and Draper, 2010; Tong et al., 2010; Tzung et al., 2015; Wang et al., 2007); however, how cysts form has not been described yet.

To investigate cystogenesis, we labeled wild-type GCs with anti-Vasa antibody between 7 and 14 days (d) (Figure 1). At 7d to 10d, Vasa positive cells were detected in wild-type as individual GCs or clustered GCs (Figure 1A, G). Individual GCs were dispersed within the gonad, while clustered GCs formed closely associated groups of cells (Figure 1B,G). At this stage, the GC nuclei had condensed DNA as revealed by DAPI with no visible nucleolus, and the nuclear cytoplasm interface was highly convoluted with a high nucleus to cytoplasm ratio (Figure 1C,G). The first somatic gonadal cells have been reported to be adjacent to the GCs at around 5d (Braat et al., 1999b; Draper et al., 2007; Leerberg et al., 2017). Similarly, we observed a few presumptive gonadal somatic cells surrounding the Vasa antibody labeled GC during this period (Figure 1). Following clustering, GCs underwent a morphological transition, such that individual GCs became difficult to discern within a Vasa positive mass (Figure 1C). This step was characterized by complex cytoplasmic and nuclear rearrangements. The most prominent feature was the appearance of a compact gonad such that individual GC boundaries were not apparent within the irregular cytoplasmic compartments devoid of Vasa and DAPI (Figure 1C). During this period, DAPI signal was reduced and Vasa distribution shifted from the perinuclear aggregates typical of PGCs to diffusely cytoplasmic, suggesting that these morphological changes may correspond to mitotic divisions (Tong et al., 2010). The simultaneous obscuring of individual cell boundaries as the multiple domains devoid of Vasa emerge suggests that the phenomena is synchronous (Figure 1C). Because this phenomenon likely corresponds to amplification of GC numbers that precedes emergence of germline cysts; hereafter, we describe these Vasa positive masses as cystogenic cells and this morphogenetic process as cystogenesis. The amplification step is characterized by cystogenic cells with a compact round nucleus and patent perinuclear Vasa granules circumscribed by scant cytoplasm (Figure 1 D,G). At 10-12d (n=7 gonads), early germline cysts were apparent. Cyst cells had compact nuclei with a prominent nucleolus and perinuclear Vasa aggregates (Figure 1 E, G). Cysts appeared to have a shared cytoplasm, but it was unclear if cyst cells were interconnected by cytoplasmic bridges such as those observed in juvenile ovaries (Marlow and Mullins, 2008). Gonadal somatic cells continued to accumulate around the GCs (Figure 1 E). Among the wild-type gonads examined at 10-12d, only one individual had Vasa positive cells with the earlier single clustered GC morphology, while the others had established germline cysts. Between 12-14d, wild-type fish had pre-meiotic germline cysts with round nuclei and a prominent nucleolus (Figure 1 F,G). At this stage, Vasa protein remained concentrated in perinuclear aggregates and diffusely cytoplasmic. The nuclei of somatic gonadal cells adopted an elongated shape as the cyst grew (Figure 1 F).

**Figure 1.**
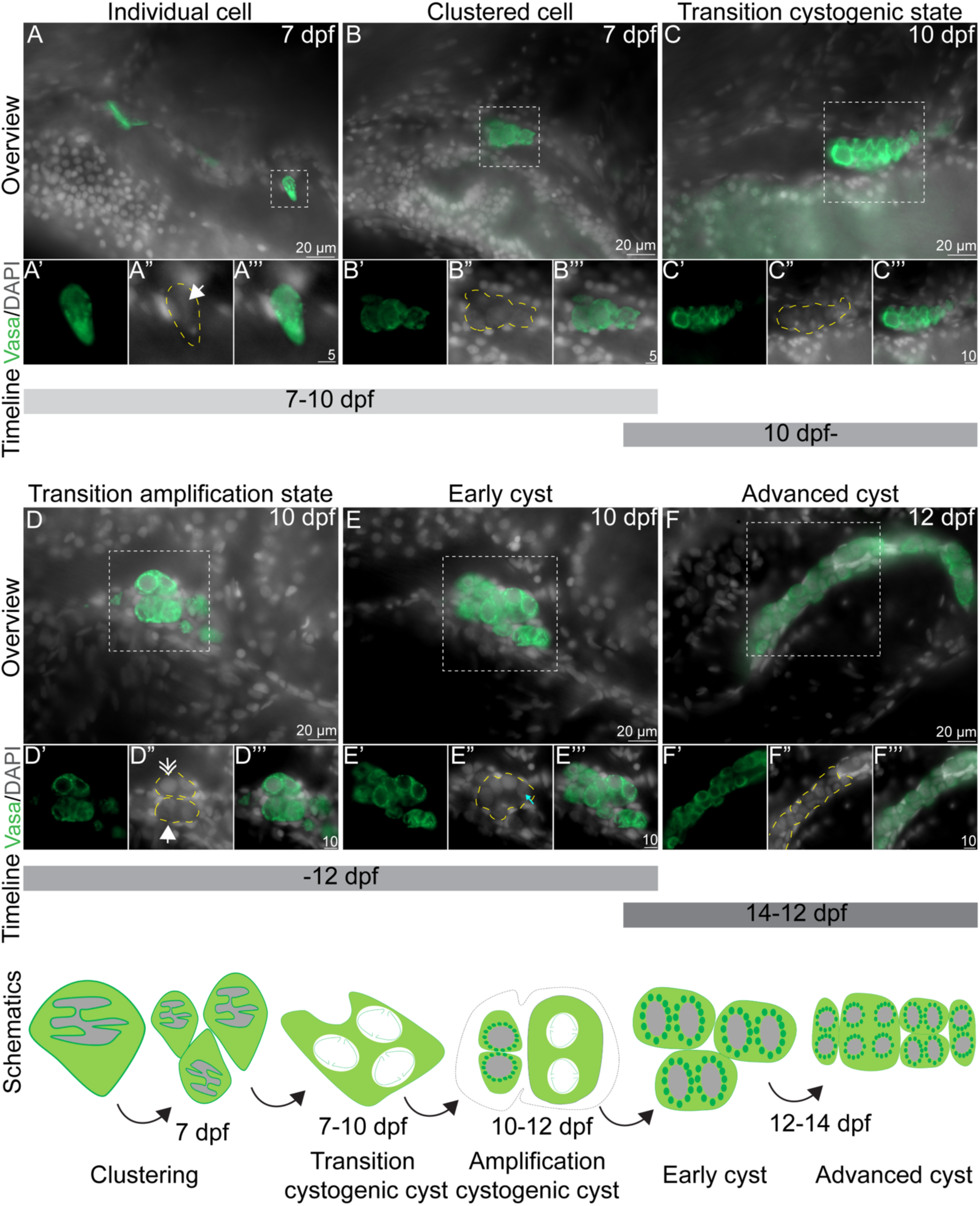
Overview of zebrafish cystogenesis. GCs are labeled with anti-Vasa antibody (Green), and nuclei (gray) are marked with DAPI. Merged overviews and magnified insets showing each channel (Vasa, in green; DAPI, in grey; merge in green/grey). Yellow dotted line delineates GC cytoplasm. (A-A’’’) GCs are individualized in the germinal epithelium and display compact DNA structures indicated by a white arrow. (B-B’’’) Individual GCs form a cluster of cells with condensed nuclei. Clustering is observed between 7-10d. (C-C’’’) GCs transit to a cystogenic state marked by compacted germ cells with irregular cytoplasmic and nuclear boundaries and diminished DAPI staining. (D-D’’’) Cell amplification by division (white filled arrowhead) and formation of germline cyst cells with round nuclei (two-chevron arrowheads), perinuclear cytoplasm becomes round with Vasa+ granules. (E-E’’’) Early cyst cells emerge with a clearly-defined cytoplasm, germ granules, and nuclei with an apparent nucleolus, visible as a dark sphere within the nucleus (aqua arrow). The transient-amplification process occurs between 10-12d. (F-F’’’) As cyst formation progresses, cystoblast cells have a well-defined cytoplasm with perinuclear granules, and prominent nucleoli. Scale bar, 10 µm. Inset 5 to 10 µm. (G) Schematics representing the cystogenesis process from individual cells that cluster, then divide, to form a premeiotic cyst.

To further examine cyst architecture, determine cyst size, and investigate whether early cysts were interconnected by cytoplasmic bridges, we labeled GCs with Vasa antibody and F-Actin with Phalloidin to visualize cell membranes and the cytoskeleton (Supplemental Figure 4). At 10d wild-type gonads (Figure 2A, B-B”) and GCs in 2-cell (Figure 2A, C-C”) and 4-cell (Figure 2A, D-D”) configurations. Cyst cells were connected by prominent ring canal structures (Figure 2E-E’). These structures were present before the reported onset of meiotic marker expression in zebrafish (Beer and Draper, 2013; Rodriguez-Mari et al., 2013; Tzung et al., 2015); therefore, we conclude that these are premeiotic GCs cysts.

**Figure 2.**
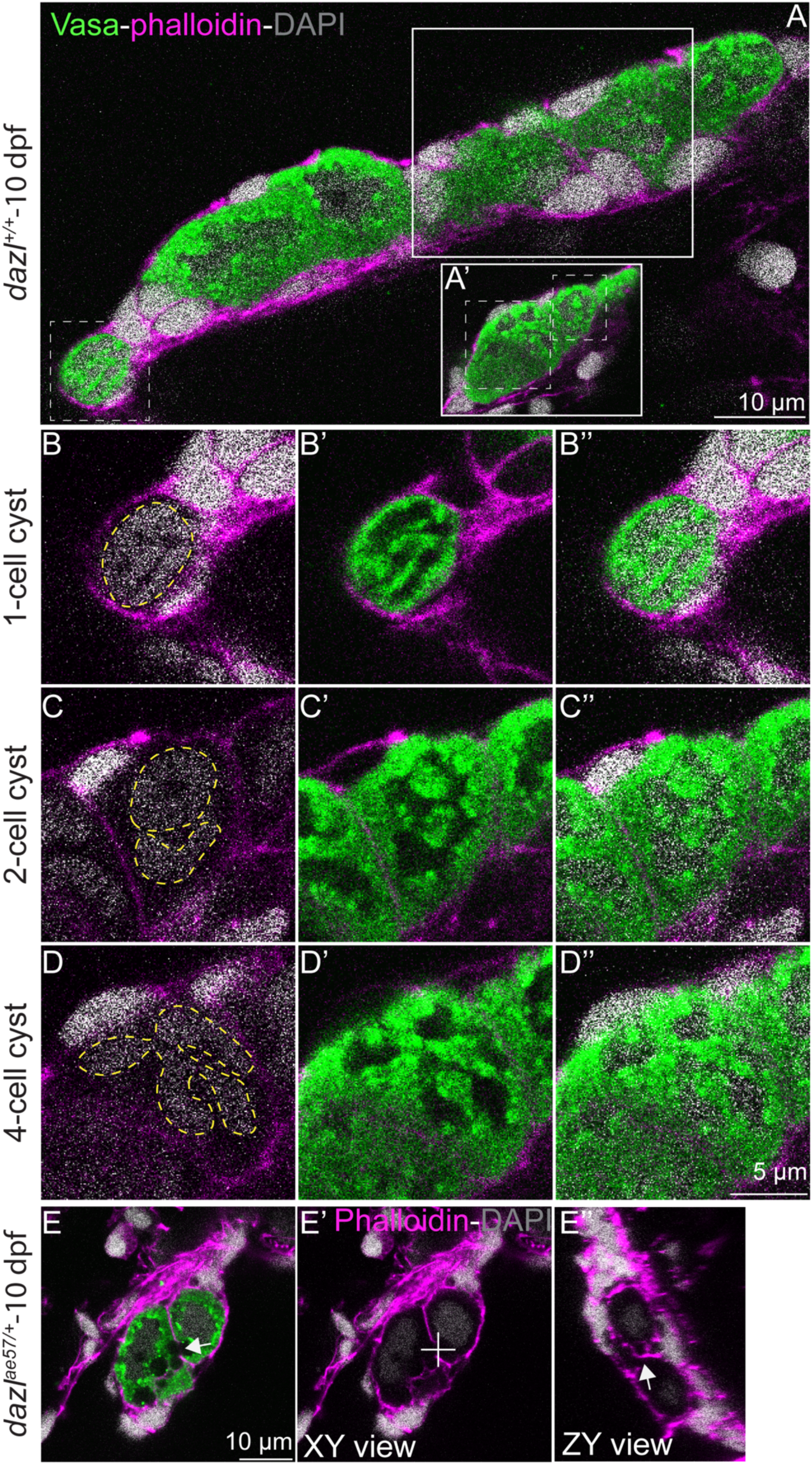
Cyst and ring canal formation in zebrafish. (A-A’) Single confocal plane of a larval gonad at 10d with nuclei labeled with DAPI (gray), GCs marked with Vasa (green), and F-actin with phalloidin (pink). Each cystogenesis stage is boxed with white dotted lines. Box with solid white line delineates the location the focal plane showing the 2 or 4 cell stage cysts. (B-B’’) 1-cell stage marked by a compact DNA (B), a high cytoplasm/nucleus ratio (B’). (C-C’’) Division produces a 2-cell stage cyst with two nuclei (yellow dotted line) surrounded by perinuclear Vasa. (D-D’’) A second round of division generates a 4-cell cyst with four nuclei (yellow dotted line). (E-E’’) Intercellular bridges in 10d larval gonad indicated by the white arrow. (E’) XY view without Vasa (green). A white cross mark the site of the orthogonal view (ZY) seen in E’’. (E’’) ZY view showing the ring (white arrow). Scale bar, 10µm or 5µm as indicated.

### dazl mutants

Although Dazl has not been shown to be required for cyst formation, we reasoned that Dazl was a compelling candidate regulator of this conserved process, since in mouse and human fetal ovary it interacts with Tex14, a regulator of ring canal formation (Greenbaum et al., 2006; Reynolds et al., 2005; Rosario et al., 2017; Zagore et al., 2018). In zebrafish *dazl* is expressed at all stages of GC development, but its function at this particular stage of germline development is unknown (Hashimoto et al., 2004; Howley and Ho, 2000; Kosaka et al., 2007; Maegawa et al., 1999). To test the hypothesis that *dazl* regulates germline cyst formation, we generated three mutant alleles disrupting zebrafish *dazl* using zinc finger nucleases (Foley et al., 2009a) and Crispr-Cas9 (Gagnon et al., 2014). We recovered *dazl^Δ7^* using zinc finger nucleases to target the first exon of *dazl* (Figure 3A). Sequencing of the genomic and mutant cDNA revealed a frameshift mutation that caused a premature stop codon within *dazl* that removes all functional domains (Figure 3B, C). Using Crispr-Cas9 methods to target exon 6 of *dazl,* we recovered two additional alleles. Sequencing of genomic DNA and mutant cDNA revealed two distinct insertion-deletion alleles (Figure 3B, C; Supplemental Figure 1). The *dazl^ae57^* allele harbors a 2bp deletion, a 9bp insertion, and 2bp substitution that results in a premature stop codon predicted to truncate Dazl and eliminate the Daz motif (Figure 3B, C). The *dazl^ae34^* allele, a 15bp insertion with a 12bp deletion resulted in a four amino acid in-frame deletion generated a truncated protein of 225 amino acids with intact RRM and Daz motifs. Genotyping assays were developed for each *dazl* mutant allele (Figure 3D; Supplemental Figure 2).

**Figure 3.**
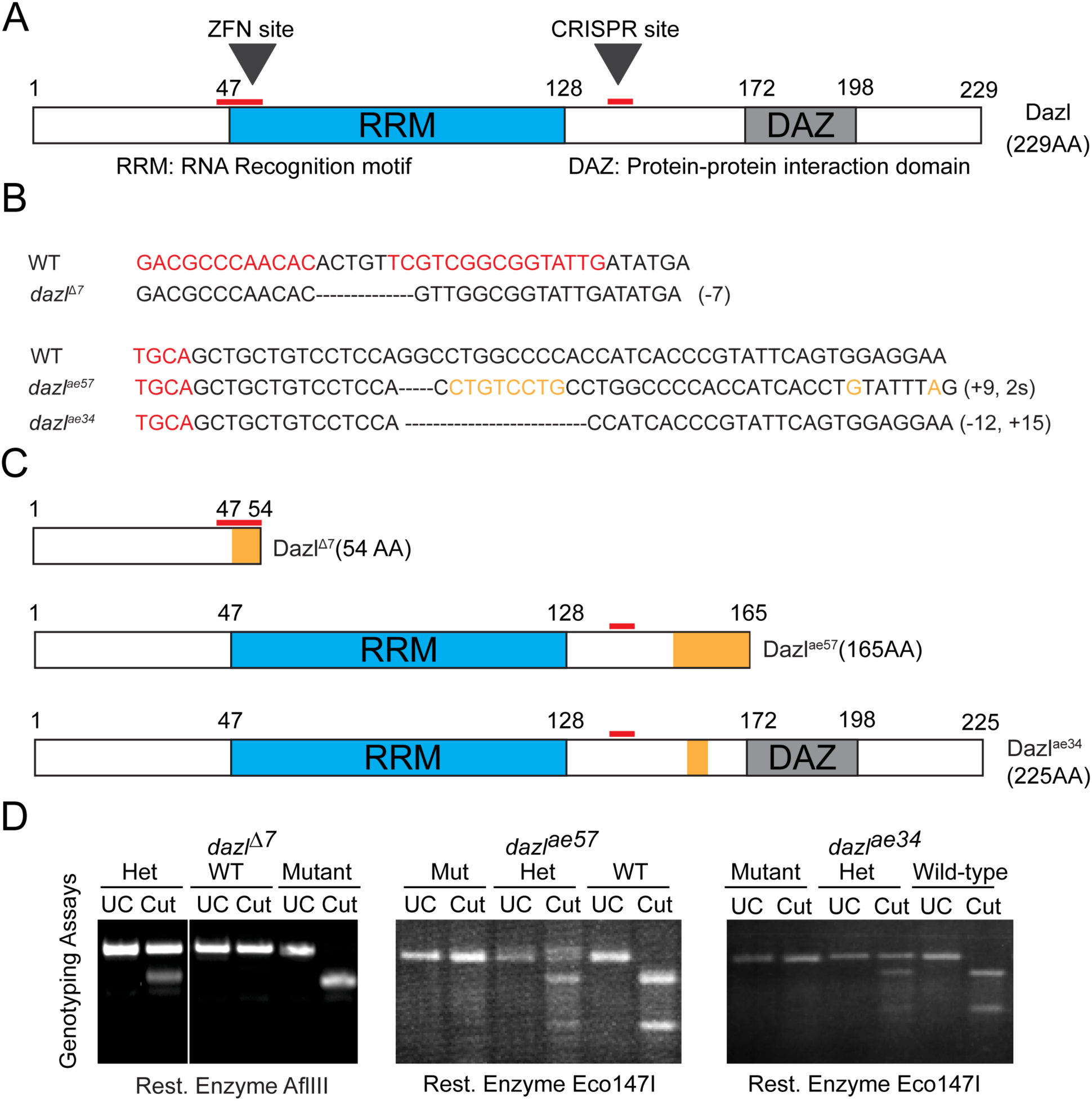
*dazl* mutagenesis strategy and fertility phenotypes. (A) Dazl protein diagram illustrating the RRM (RNA Recognition Motif) and DAZ domains with their respective amino acid regions. The ZFN and CRISPR sites are indicated above by a red line. (B) Both partial zinc finger nucleases and Cas9 binding sites are highlighted in red. *dazl* alleles generated by ZFN (*dazl^Δ7^)* and CRISPR (*dazl^ae57^*and *dazl^ae34^).* Deletions are represented by a dashed line and the substitution is highlighted in orange. (C) Diagram of alleles generated. 1) ZFN allele causes a 7 bp deletion and leads to a premature stop codon that eliminates all functional domains. 2) *dazl^ae57^,* a CRISPR-Cas9 induced allele creates a 9bp insertion and 2bp substitution leading to a premature stop codon just after the RRM domain. The third allele, also obtained with CRISPR-cas9 mutagenesis, induced a 12bp deletion and 15bp insertion causing an in-frame deletion between the RRM and DAZ domains. (D) Representative PCR based DCAPS genotyping assay for each allele. Heterozygotes are composed of the wild-type allele (upper band) and the mutant allele (lower band) in the case of *dazl^Δ7^*. *dazl^ae57^* or *dazl^ae34^* heterozygotes contains the wild-type allele (lower band) and the mutant allele (upper band).

### Zygotic dazl is dispensable for PGC specification and migration

First, we determined whether GCs were specified in *dazl* mutants by examining the GC marker *nanos3* (Koprunner et al., 2001) at shield stage and found no differences in PGC number, position or morphology between *dazl* mutants and siblings (Figure 4A; n=45 embryos; 12 *dazl*^+/+^; 22 *dazl*^ae57/+^; 11 *dazl^ae57/ae57^*). Having confirmed that GCs were specified, we examined two GC markers, *nanos3* (Koprunner et al., 2001) (Figure 4B) and Vasa, and quantified GC number and position at 30hpf. GCs were present in all progeny of *dazl* heterozygous parents at this stage (Figure 4C-J). Although the number of GCs in individuals varied, no significant differences in GC number were found at this stage (Figure 4J). Moreover, Vasa labeled germ granules of PGCs were indistinguishable between *dazl* mutants and siblings (Figure 4D, F, H insets; n=28; 5 *dazl*^+/+^; 17 *dazl*^ae57/+^; 6 *dazl^ae57/ae57^*). Defective GC migration can lead to death of GCs or loss of germline identity (Draper et al., 2007; Gross-Thebing et al., 2017; Weidinger et al., 2003), and *dazl* has been implicated in PGC migration in *Xenopus* (Houston and King, 2000). Therefore, we examined Vasa labeled PGC position in *dazl* mutants. Vasa positive PGCs were present and indistinguishable between *dazl^ae57^* and *dazl^Δ7^* mutants and siblings. Although the occasional stray germ cell was observed, there were no significant differences in PGC migration between *dazl* mutants and siblings (Figure 4B-H). Moreover, we observed no differences between *dazl^ae57/ae57^, dazl^Δ7/Δ7^;* or *dazl^ae57/Δ7^* mutants in any of the assays performed in this study. Therefore, we conclude that if *dazl* is required for GC specification, PGC granule formation, or germ cell viability, maternal *dazl* must be sufficient at these stages.

**Figure 4.**
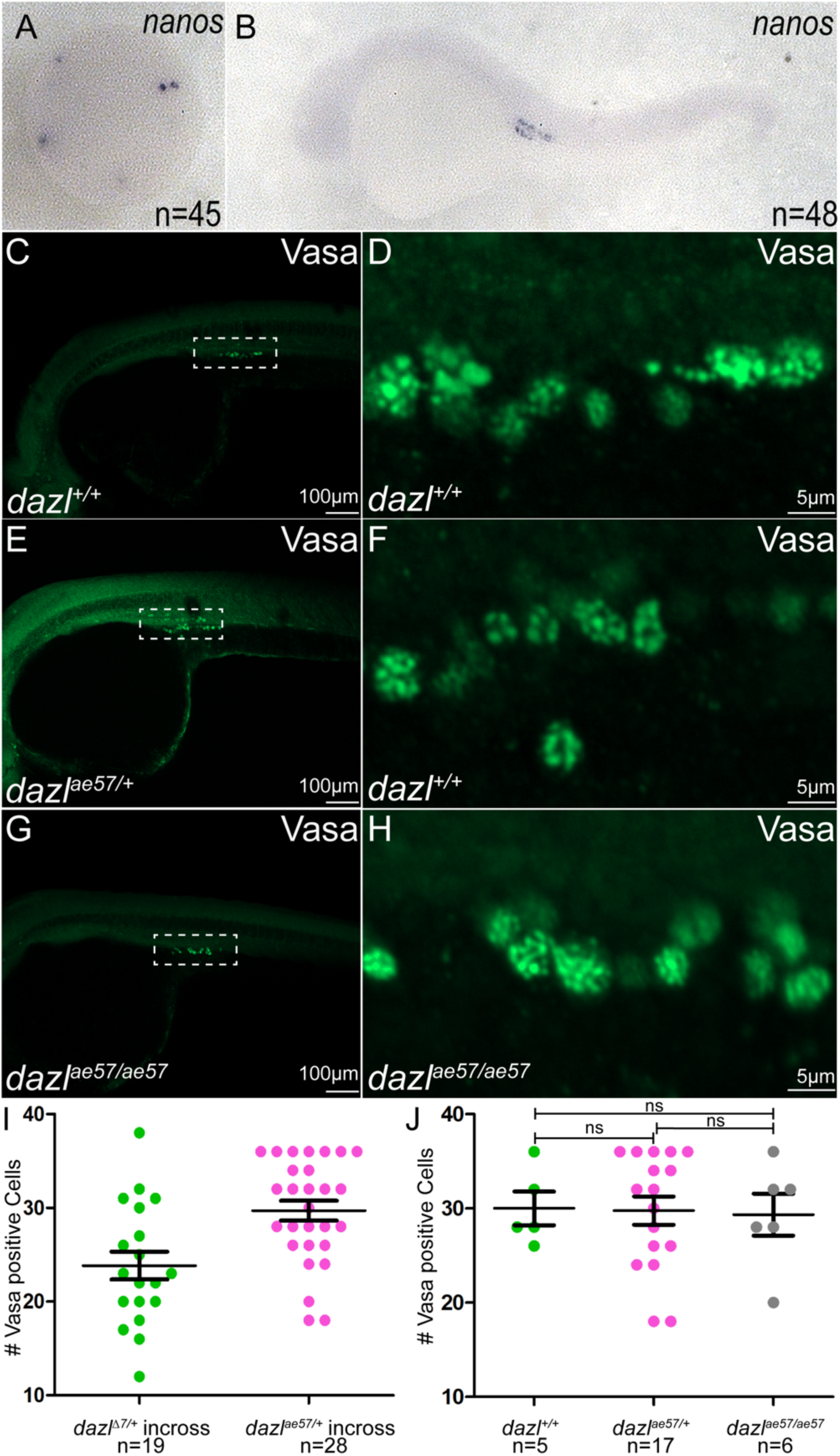
Zygotic *dazl* is dispensable for PGC specification and migration. (A, B) *In situ* of *nanos3* RNA in representative progeny of *dazl^ae57/+^* intercrosses *nanos3* at sphere (A) stage (animal pole view) and 30 hpf (B; lateral view). (C, H) Vasa immunostaining of *dazl^+/+^* (C, D), *dazl^ae57/+^* (E, F) and *dazl^ae57/ae57^* (G, H) embryos at 30hpf. A magnified view of the GC region of each genotype is shown in each inset reveals no difference in germ granules (D, F, H).. Dorsal is oriented toward the top in panels (B-H). (I) Quantification of Vasa+ GCs in progeny of a *dazl^Δ7/+^* (left) or *dazl^ae57/+^* heterozygote intercross. (J) Quantification of Vasa+ cells in a *dazl^ae57/+^* heterozygote intercross grouped according to genotype (*dazl^+/+^*; *dazl^ae57/+^*; *dazl^ae57/ae57^*). Statistics were performed between *dazl^+/+^* and *dazl^ae57/+^* (ns:0.9375), *dazl^+/+^* and *dazl^ae57/ae57^*(ns:0.8261) *dazl^ae57/+^* and *dazl^ae57/ae57^* (ns:0.8818).

### dazl is required for cystogenesis

To determine if *dazl* was required for cystogenesis we examined Vasa and actin labeled with Phalloidin in *dazl* mutants. Between 7-10 d, Vasa persisted in *dazl* mutants, indicating zygotic Dazl is not required for early Vasa expression (Figure 5; Supplemental Figure 3). As in wildtype (Figure 5A; Supplemental Figures 3, 4, 5), Vasa positive cells of *dazl* mutants, were found as single and clustered individuals (Figure 5B; Supplemental Figures 3, 4, 5). Somatic gonadal cells were similarly apparent next to *dazl* mutant GCs (Figure 5; Supplemental Figure 3). Like wildtype at 10d (Figure 5C; Supplemental Figures 3 and 4), *dazl* mutant cells transitioned to a cystogenic state and underwent a synchronous nuclear/cytoplasmic rearrangement with DAPI becoming faint (Figure 5D; Supplemental Figures 3 and 4). Some dazl mutant GCs showed evidence of amplification, including a compact nucleus (DAPI), but Vasa was diffusely cytoplasmic, and Vasa perinuclear aggregates were less apparent at this time (Supplemental Figures 3, 4, 5). Taken together these results indicate that initial transition and early amplification phases do not require *dazl*.

**Figure 5.**
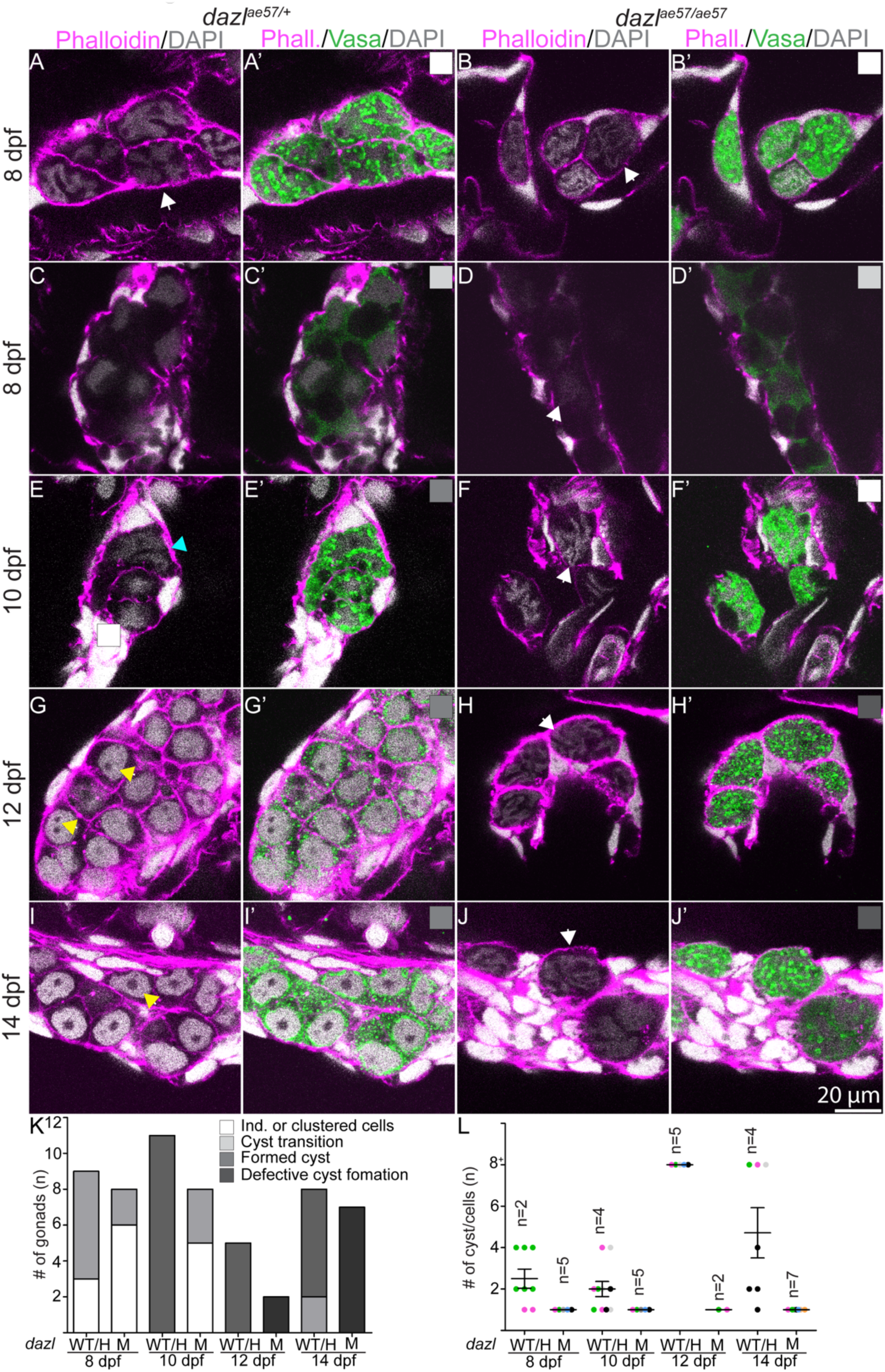
Cystogenesis in zebrafish. Larval gonads labeled with Vasa (green), DAPI, and F-actin (fluorescent conjugated phalloidin). (A-A’-C-C’-E-E’-G-G’, I-I’) Single confocal plane of *dazl^ae57/+^* gonad showing cystogenesis between 8-14d. (A-A’-C-C’-E-E’) Clustered Vasa+ individuals (white arrowheads) exhibit the first step of cystogenesis with compaction and compartmentalization into irregular nuclear, Vasa+ and Vasa-domains (C-C’-D-D’) at 8-10d. Although initial compaction and compartmentalization occurs in both wild-type and mutant genotypes, (E-E’) GCs of wild-type genotype in resolve as cysts with multiple nuclei evident (Aqua arrowhead), (F-F’) whereas *dazl* mutants return to the pretransition morphology. (G-G’) Amplification of GC numbers, round shaped nuclei with one to two nucleoli (yellow arrowheads) at 12d (n=3). (I-I’) Cells are organized in cysts with pre-meiotic nuclei with a prominent nucleolus (yellow arrowhead). *dazl^ae57/+^* (8 d, n=3 (A-A’); 8d, n=6 (C-C’); 10d, n=11; 12d, n=5; 14d, n=6). (B-B’-D-D’-F-F’-H-H’, J-J’) age-matched *dazl^ae57/ae57^* gonads. At each stage after the cyst transition, *dazl^ae57/ae57^* germ cells return to the morphology of individuals with convoluted DNA (white arrows) and lack cyst organization. *dazl^ae57/ae57^* (8 d, n=6 (B-B’); 8d, n=2 (D-D’); 10d, n=5; 12d, n=2; 14d, n=7). Number of samples (n) are indicated in each panel. Scale bar, 20µm. (K) Quantification of each category between 8-14d in *dazl^ae57/+^* and *dazl^ae57/ae57^* gonads. Note that each category is displayed with the corresponding box on the top right of each sample. (L) Quantification of cells per cyst at 8-14d in *dazl^ae57/+^* and *dazl^ae57/ae57^* gonads.

Next, we examined *dazl* mutants at 12d, when cysts emerge in wildtype. In contrast to the interconnected Vasa positive cyst cells in wild-type and *dazl* heterozygotes (Figure 5E), organized cysts did not form in *dazl* mutants, instead the *dazl* mutant cells reverted to the earlier intermingled individual cell morphology (Figures 5F; Supplemental Figure 3). The cytoplasm of the mutant cells appeared convoluted in appearance, and perinuclear enrichment of Vasa granules was not evident (Supplemental Figure 3). At 14d in contrast to wild-type gonads which were full of cysts (Figure 5G, I; Supplemental Figures 3 and 4), *dazl* mutant GCs remained as individuals indicating a failure of or severe delay in cystogenesis (Figure 5H, J; Supplemental Figures 3, 4, 5). Because WT cells appear to become smaller as a result of cystogenesis, we measure cell size prior to and after cystogenesis transition. Using 3D imaging software, we determined cell area and volume at 8-14d. *dazl^ae57/+^* GCs decreased significantly in area and volume, indicating the divisions are reductive in nature (Supplemental Figure 6). In contrast, *dazl* mutant cell area and volume did not decrease, suggesting failed division (Supplemental Figure 6). Because somatic gonadal cells encapsulating the GCs were intact, and because *dazl* expression is GC specific (Hashimoto et al., 2004; Maegawa et al., 1999), Dazl likely acts within the GCs to promote cystogenesis.

Failure to generate a cyst in *dazl* mutants could be due to abnormal cyst architecture or cytokinesis such as failure to arrest the cytokinetic furrow, particularly given that Dazl interacts with cytokinesis and ring canal factors (Reynolds et al., 2005; Rosario et al., 2019; Rosario et al., 2017; Zagore et al., 2018). Actin is a prominent marker of intercellular connections and subcellular structures of GC cysts in other species, including the spectrosome of GSCs, and the branched fusome that connects germline cyst cells (de Cuevas et al., 1997; Kloc et al., 2004; Lin and Spradling, 1995; Snapp et al., 2004; Spradling et al., 1997). Close inspection of actin (labeled with Phalloidin) in gonads with wild-type genotypes at 8 and 10d revealed an actin rich density within GCs, potentially a spectrosome (Figure 6A-A’; C-C’). These aggregates were also present in *dazl* mutant GCs at 8 and 10d (Figure 6B-B’; D-D’). By 12d, when cysts are abundant in wild-type, actin was present in branched structures reminiscent of the fusome of other species (Hime et al., 1996; Kloc et al., 2004; Warn et al., 1985) (Figure 6E-E’). In contrast, at this stage in *dazl* mutants the spectrosome-like aggregates persist in Vasa+ cells (Figure 6F-F’). By 14dpf these actin structures resolved and were no longer detected in wild-type cysts (Figure 6G-G’, I). However, the actin densities persisted and appeared to have duplicated as two actin aggregates were present in *dazl* mutant GCs as opposed to the single aggregate observed at 8 and 10d in *dazl* mutants (compare Figure 6H-H’ and 6B-B’, J), possibly reflecting failed division or cell fusion.

**Figure 6.**
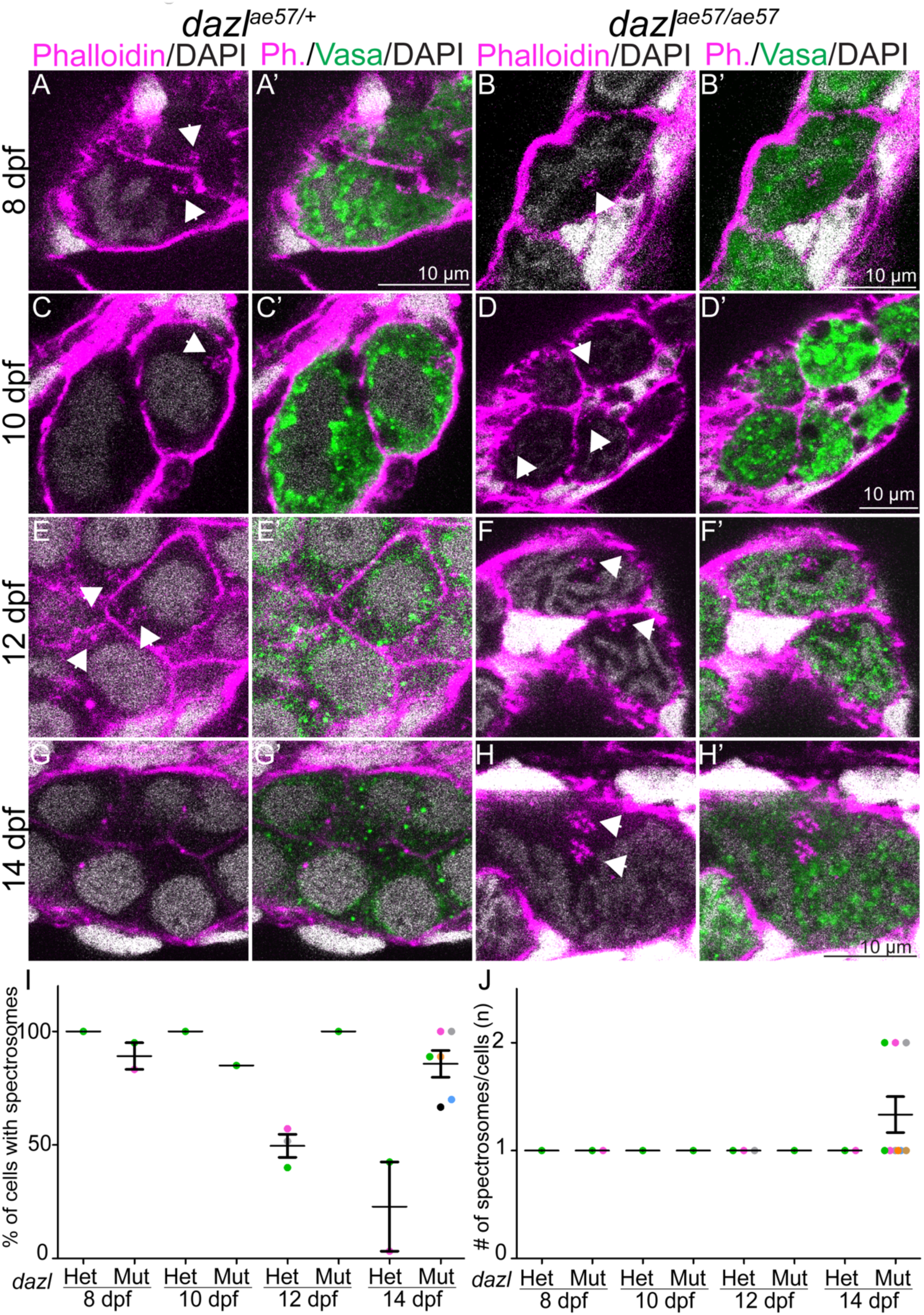
*dazl* mutant cells retain spectrosomes. Single confocal plane of *dazl^ae57/+^* or *dazl^ae57/ae57^* gonad between 8-14d labeled with Vasa and fluorescent conjugated phalloidin which marks spectrosomes and nuclei labeled with DAPI. (A-A’, C-C’, E-E’, G-G’) The spectrosome indicated by the white arrow is present transiently between 8-12d in *dazl^ae57/+^* gonads. (B-B’, D-D’, F-F’, H-H’) Spectrosome structures (white arrow) are present in *dazl* mutant cells between 8-14d. (H-H’) At 14d, double spectrosomes are observed. Scale bar, 10µm. (I) Quantification of the number of cells containing a spectrosome within each gonad. (J) Quantification of spectrosomes per cell.

The main structures connecting cells within cysts are the ring canals, thought to be the products of arrested cytokinesis (Haglund et al., 2011; Hime et al., 1996; Robinson and Cooley, 1996). At 10d ring canals were present between cyst cells in wild-type (Figure 7A-A”) and in *dazl* mutants (Figure 7B-B”), consistent with normal onset of cystogenesis in the absence of *dazl.* In wild-type cysts conspicuous ring canals remained at 12-14d (Figure 7C-C”; E-E”). In contrast, ring canals appeared collapsed in *dazl^-/-^* with membranes closely abutting one another rather than in an open ring configuration as cells individualized (Figure 7D-D”; F-F”). Taken all together, we conclude that *dazl* is essential for incomplete cytokinesis/type-II divisions and germline cyst formation.

**Figure 7.**
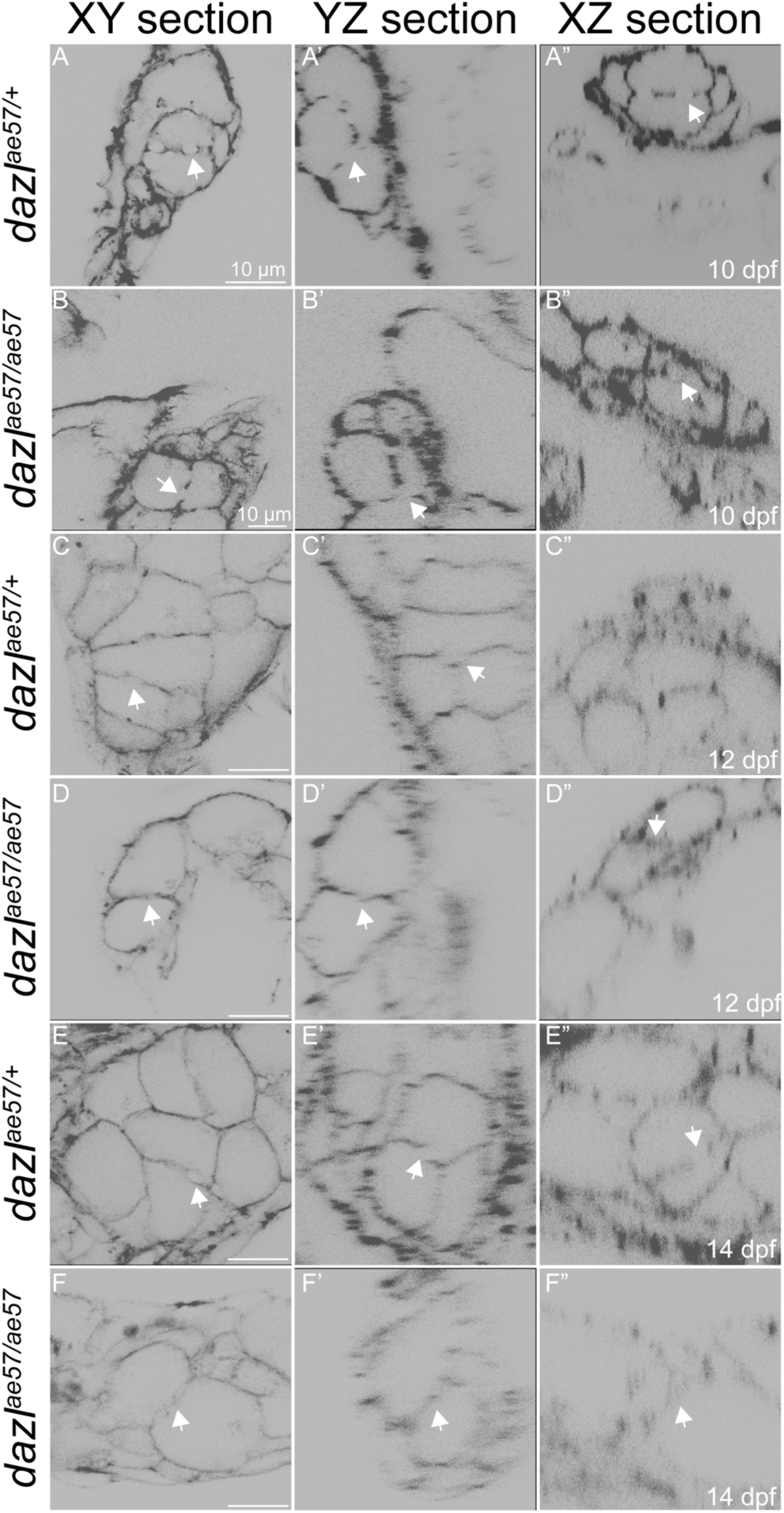
Zygotic *dazl* mutant cells do not maintain ring canals. Single confocal plane of representative *dazl^ae57/+^* or *dazl^ae57/ae57^* gonads between 10-14d labeled with fluorescent conjugated phalloidin (Actin) depicting the ring canal. Each stage is represented with a sagittal view (XY) or orthogonal views (YZ or XZ) of a Z-stack reconstruction. (A-A’-A’’, C-C’-C’’, E-E’-E’’) Ring canals at each stage between 8-14d indicated by a white arrowhead in *dazl^ae57/+^* gonads. Note the circular shape. Number of gonads analyzed: 10 d, n=3 (A-A’-A’’); 12d, n=3 (C-C’-C’’); 14d, n=3 (E-E’-E’’). (B-B’-B’’, D-D’-D’’, F-F’-F’’) Presence of ring canals at 10d in *dazl^ae57/ae57^* cells. At 12 and 14d, ring canals are not maintained (white arrows). Number of gonads analyzed: 10 d, n=4 (B-B’-B’’); 12d, n=2 (D-D’-D’’); 14d, n=3 (F-F’-F’’). Scale bar, 10 µm.

### Type-II/cystogenic divisions are required for fertility

As discussed, two modes of GSC divisions, type-I/direct differentiating and type-II/cystogenic, have been described (Marlow and Mullins, 2008; Nakamura et al., 2010; Saito et al., 2007). In mouse, the ring canals that connect cyst cells are dispensable for fertility in females but are required in males (Greenbaum et al., 2006). In contrast, defects in ring canal formation in *Drosophila* disrupt fertility of both sexes (Hime et al., 1996; Yue and Spradling, 1992). Type-II divisions fail in *dazl* mutants. To determine if cystogenic divisions are required for fertility in zebrafish, *dazl* mutants were raised to adulthood. As expected based on the germline specific expression of *dazl* (Kosaka et al., 2007; Maegawa et al., 1999), no overt morphological defects were observed among *dazl^-/-^*. Although mutants were viable to adulthood, *dazl^Δ7/Δ7^* (n=11) and *dazl^ae57/ae57^* (n=6) adults were exclusively sterile males. In contrast, *dazl^ae34/ae34^* were fertile (n=23 males; n=6 females).

To confirm that infertility of *dazl* mutants was specific for *dazl* mutation and to assess mutant allele strengths we performed complementation tests. While heterozygosity for each allele caused no phenotypes or fertility deficits (Figure 8A, B, D’, D’-E, E’ n=9), *dazl^ae57/Δ7^* mutants were exclusively sterile males (Figure 8C, F, F’ n=2). Examination of Vasa-labeled GCs (Leerberg et al., 2017) and DNA with DAPI, revealed oocytes (Figure 8G) or sperm (Figure 2H) in wild-type siblings at 2 months of age, and lack of Vasa+ GCs in *dazl^ae57/Δ7^* mutants (Figure 9I; n=2). In contrast, *dazl^ae57/ae34^* mutants were fertile males or females, albeit with a male bias (n= 5 females, n=10 males). Based on these observations, we conclude that *dazl* mediated cystogenesis/type-II division is essential for GSC establishment and fertility in zebrafish. Moreover, *dazl^ae57^* and *dazl^Δ7^* are strong loss of function alleles, but *dazl^ae34^* retains sufficient function to support normal germline development.

**Figure 8.**
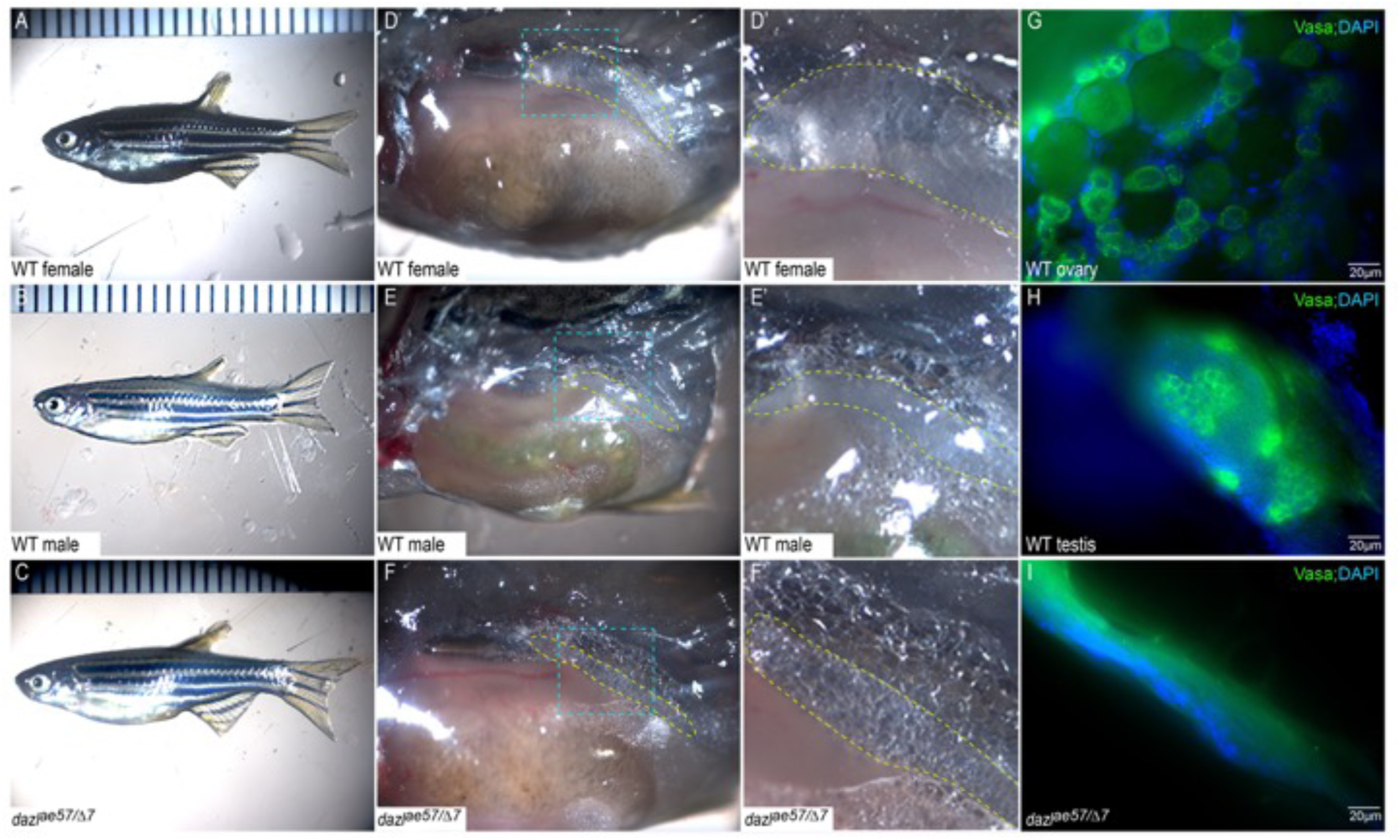
Zygotic *dazl* mutants are sterile males. Overall morphology of representative adult wild-type female (A), male (B), *dazl^Δ7/ae57^* compound heterozygote (C). Dissected trunks (D-F). The gonad region is outlined in yellow for the wild-type female (D), male (E), and *dazl^Δ7/ae57^* compound heterozygote (F). Higher magnification views of region outlined in blue (D’-F’). Oocytes in female gonads (D’), and sperm/ testicular structures of males (E’). No identifiable gametes or GCs were detected in compound heterozygotes (F’). Gonads labeled with Vasa reveal the oocytes in (G), spermatocytes (H), and a gonad devoid of GCs (I).

**Figure 9.**
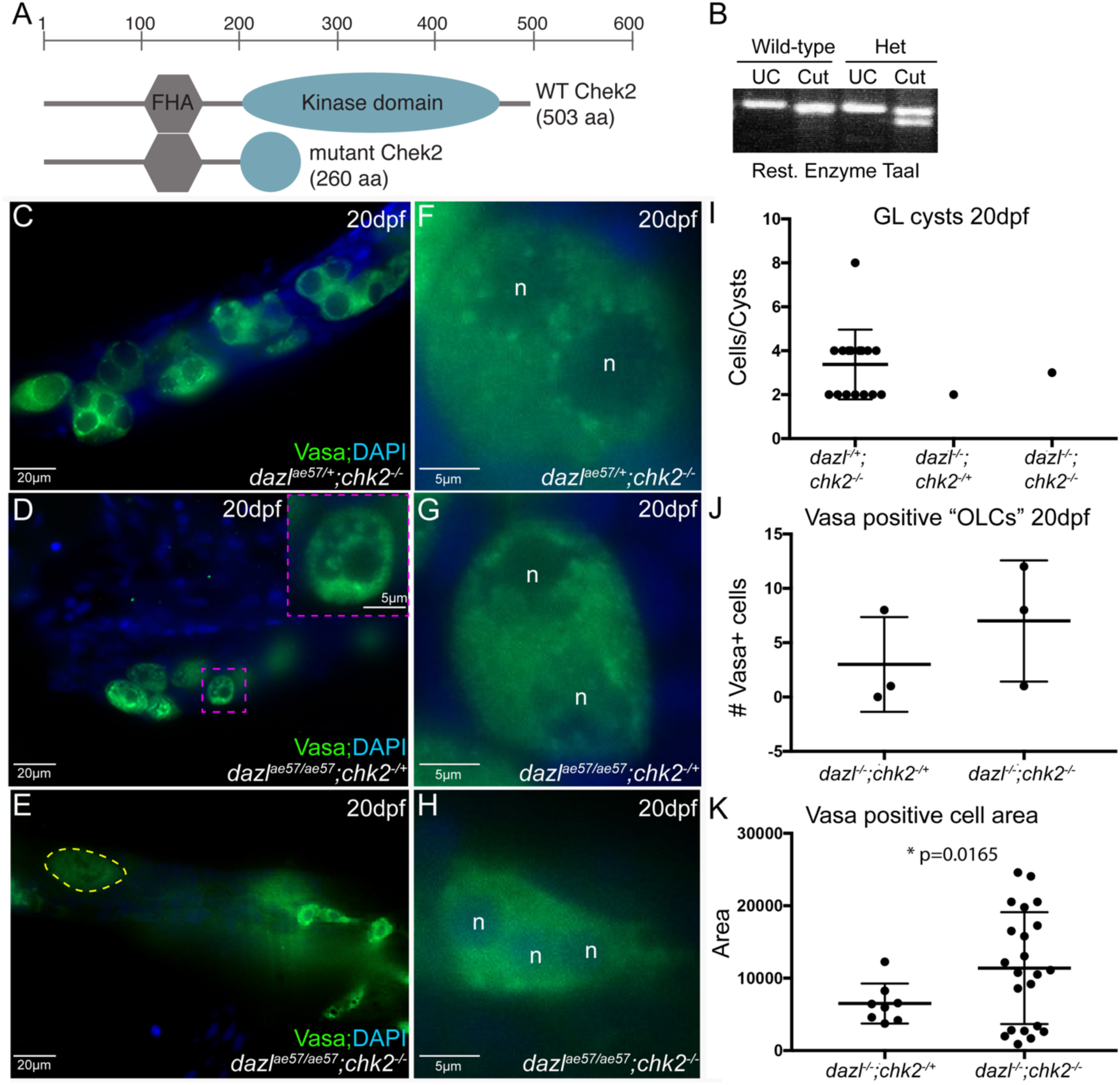
*chk2* inactivation does not prevent GC loss. (A) Schematic of wild-type and mutant Chk2 proteins. (B) Representative genotyping assay. TaaI cuts the mutant allele. (C-D) GC clusters of *dazl^ae57/+^;chk2^sa20350/sa20350^*. Presence of Vasa granules surrounding the GC nucleus. (D-G) Vasa+ GCs of *dazl^ae57/ae57^*; *chk2^sa20350/+^*. Note the vacuolization of the cell enlarged in the inset (E-H) Vasa+ GCs of *dazl ^ae57/ae57^*; *chk2^sa20350/sa20350^*. (I) Quantification of germline cysts for *dazl ^ae57/+^*; *chk2^sa20350/+^*, dazl^ae57/ae57^; chk2^sa20350/+^ and *dazl ^ae57/ae57^*; *chk2^sa20350/+^* (n=3 gonads/genotype). (J) Quantification of large Vasa+ GCs of *dazl^ae57/ae57^*; *chk2^sa20350/+^* and *dazl^ae57/ae57^*; *chk2^sa20350/sa20350^*. (K) Quantification of the _cell area in Vasa+ GCs in *dazl*_*ae57/ae57*_; *chk2*_*sa20350/+* _and *dazl*_*ae57/ae57_; chk2_sa20350/sa20350*. Scale bars, 20 µm (C-E) and 5µm (F-H).

### Checkpoint inactivation cannot suppress loss of germ cells in dazl mutants

Loss of germline has been reported to occur either by cell death or failure to maintain GC fate and differentiation as somatic cell types (Gross-Thebing et al., 2017; Kossack and Draper, 2019; Leerberg et al., 2017; Rodriguez-Mari et al., 2010; Slanchev et al., 2005; Uchida et al., 2002; Wang et al., 2007). During normal development, apoptosis of GCs is associated with male-specific differentiation (Pradhan et al., 2012; Rodriguez-Mari et al., 2010; Siegfried and Nusslein-Volhard, 2008; Slanchev et al., 2005; Tzung et al., 2015; Uchida et al., 2002; Wang et al., 2007), and mutation of the apoptosis regulator, *tp53,* can suppress cell death and loss of oocytes in several zebrafish mutants, including *zar1, brca2,* and *fancl* (Miao et al., 2017; Rodriguez-Mari et al., 2010; Shive et al., 2010). Because zebrafish *dazl* mutant GCs are not detected after 18d, we investigated whether blocking cell death by *tp53* checkpoint activation could suppress germline loss in *dazl* mutants. To do so, we crossed the *tp53* mutation into the *dazl^ae57^* background to generate double mutants. Analysis of dissected adult (>6 months) gonads revealed ovary or testis in *tp53* mutants that were heterozygous for *dazl^ae57^* (n=9). In contrast, all *dazl^ae57/57^* whether *tp53* heterozygous (n=4) or *tp53^-/-^* (n=5) lacked GCs and were sterile males. To investigate whether *tp53* mutation could prolong GC viability in *dazl* mutants, we analyzed Vasa-labeled GCs at 40dpf. As expected, OLCs or oocytes were present in *tp53* mutants that were heterozygous for *dazl^ae57^* (Supplemental Figure 6; n=6). In contrast, no GCs were detected at 40d in *dazl^ae57/57^* gonads regardless of *tp53* genotypes (n=6 heterozygous; n=4 *tp53^-/-^*) (Supplemental Figure 7). Therefore, we conclude that GC loss in *dazl* mutants is Tp53 independent.

In mouse, some infertility phenotypes associated with germline cell death can be suppressed by *chek2* mutation, which also acts via *tp63* (Bolcun-Filas et al., 2014). Zebrafish have a single *chek2* gene on chromosome 5. The zebrafish *chek2^sa20350^* mutant allele was recovered in the Sanger mutation screen; however, the mutant phenotype has not been reported (Kettleborough et al., 2013). We obtained the *chk2^sa20350^* nonsense allele (Kettleborough et al., 2013), which we confirmed by sequencing genomic and cDNA from mutants harboring a C to A mutation that creates a premature stop codon (Q-stop) to yield a truncated Chek2 protein lacking the protein kinase domain (Figure 9A; Supplemental Figure 8) and developed a DCAPs genotyping assay (Figure 9B). Mutation of *chek2* does not disrupt meiosis or fertility in *Drosophila* (Abdu et al., 2002) or mouse (Bolcun-Filas et al., 2014). Similarly, we found that *chek2* mutation caused no overt phenotypes, and did not interfere with viability, sexual differentiation or fertility in zebrafish as both male and female adult *chek2* mutants were fertile (n=9 females and n=13 males). Therefore, we conclude that *chek2* is not required for germline development or fertility in zebrafish.

To determine if inactivation of *chk2* could suppress germline loss in *dazl* mutants, *chek2^sa20350^* and *dazl^ae57^* double mutants were generated. The resulting progeny were examined at 20d and as adults for sex-specific traits and the presence of Vasa+ GCs. At 20d, Vasa+ GCs were present in *dazl^ae57/+;^chk2 ^sa20350/ sa20350^* mutants (Figure 9C,F), and in some *dazl^ae57/ae57^* mutants that were also *chk2* heterozygotes (Figure 9D, G) or homozygous mutants (Figure 9E,H). In contrast to *dazl^ae57/+;^chk2 ^sa20350/ sa20350^* mutants, which had numerous Vasa+ cysts ranging in size from 2-8 cells, Vasa+ cells were rarely detected in *dazl* mutants (Figure 9I). At this stage a few large vacuolated Vasa+ cells were detected in some *dazl^ae57/ae57^;chk2 ^sa20350^* heterozygotes and *dazl^ae57/ae57^;chk2 ^sa20350/ sa20350^* mutants (Figure 9B inset, J,K). The GCs of *dazl^ae57/ae57^; chk2 ^sa20350/ sa20350^* mutants were more numerous and larger compared to *dazl^ae57/ae57^;chk2^sa20350^* heterozygotes (Figure 9K), an indication that *chk2* loss may prolong GC viability; However, loss of *chk2* could not suppress eventual germline loss, as adult *dazl;chk2* mutants were phenotypically males that lacked germ cells (n=3) like their *dazl* single mutant siblings (n=4), while their *dazl* heterozygous siblings were either fertile males or females (n=11). Taken together these results indicate that *dazl* is required for cyst development and maintenance of the germline by a mechanism that acts independent of *chk2* and *Tp53* checkpoints.

## Discussion

Our study examines the earliest stages of gonadogenesis and provides evidence that the conserved RBP, Dazl, is required for germline cyst formation and plays critical roles in germ cell amplification, and acts upstream of meiosis and establishment of germline stem cells to promote fertility. Immunoaffinity screens for Dazl targets have identified regulators of incomplete cytokinesis (Kim et al., 2015; Reynolds et al., 2005; Rosario et al., 2019; Rosario et al., 2017; Zagore et al., 2018). This work provides genetic evidence that Dazl is required to form germline cysts interconnected by ring canals. Specifically, *dazl* mutant GCs initiate cyst formation, but ring canals collapse, such that mutant cells ultimately remain as individuals that fail to differentiate as meiocytes. In addition to promoting type-II cystoblast divisions we show *dazl* acts upstream of meiotic entry and GSC establishment. Finally, we show that Dazl and type-II cystogenic divisions are essential for fertility as GCs are lost from *dazl* mutants by a mechanism independent of meiotic checkpoint regulators.

### Conserved dazl functions in gametogenesis and the PGC to meiotic transition

In cultured cells, *dazl* promotes meiotic entry (Chen et al., 2014; Haston et al., 2009a; Jung et al., 2017; Yu et al., 2009). Similarly, in mouse *dazl* is required for “licensing” or acquisition of meiotic competence and to generate gametes; *dazl* mutant GCs remain PGC like and fail to develop as male or female in response to masculinizing or feminizing signals (Gill et al., 2011; Hu et al., 2015b). In medaka, *dazl* is also required for gametogenesis and GC development (Li et al., 2016; Xu et al., 2007) but is not required for sexually dimorphic gonad development in response to somatic cues (Nishimura et al., 2018). Here we show that zygotic *dazl* is required for gametogenesis and meiotic progression in zebrafish. However, in contrast to mouse *dazl* mutant gonads which fail to sexually differentiate, and medaka *dazl* mutants, which develop as either sex (Nishimura et al., 2018), zebrafish *dazl* mutants develop exclusively as males. In medaka, recovery of male or female *dazl* mutants with only early PGCs provided evidence that PGC-like cells could support development of both sexes (Nishimura et al., 2018). Although, PGCs were initially present in zebrafish *dazl* mutants, no females were recovered. Thus, in zebrafish PGCs devoid of *dazl* are not sufficient to support cystogenesis, establishment of GSCs, or a fertile gonad. In agreement with this notion, recent work in mouse and pig indicates that commitment to germline fate only occurs after the PGCs reach the gonad and requires *dazl* (Nicholls et al., 2019).

### A role for dazl in germline cyst formation

In many organisms, proliferation of GCs before meiosis is often accompanied by synchronous, incomplete divisions that form a germline cyst. During cyst formation, GSCs form cystoblasts by mitotic divisions (Leu and Draper, 2010; Pepling and Spradling, 1998). Here we characterized cystogenesis in juvenile zebrafish gonads and demonstrate a requirement for Dazl in germline cyst formation. This process begins with congregation of individual migratory GCs that then undergo synchronous division to generate premeiotic cysts. Notably, as we observe in zebrafish *dazl* mutants, *dazl^-/-^* GCs of mouse (Chen et al., 2014; Gill et al., 2011; Hu et al., 2015a; Lin et al., 2008) and medaka (Nishimura et al., 2018) remain as individual cells, which were described as PGC-like in mice and medaka (Chen et al., 2014; Gill et al., 2011; Nishimura et al., 2018). Here we show that PGCs lacking *dazl* begin the cystogenic process, including formation of ring canals. However, without Dazl, ring canals collapse signifying failure to mature bridges or completion of cytokinesis. In wild-type, the spectrosome-like actin aggregates progress to branched fusome-like structures that ultimately resolve in late cysts. It is unclear why these structures seem to disappear once the cysts form. Labeling and lineage tracing will be required to determine if cyst breakdown occurs as has been shown in mice, or if instead cysts are stable and there is selection as occurs in *Drosophila* (de Cuevas et al., 1997; Lei and Spradling, 2013; Pepling and Spradling, 1998). In contrast, actin remains in spectrosome-like aggregates, which are subsequently duplicated in *dazl* mutants. These duplicated structures potentially indicate cell fusion or more likely failed cytokinesis because cell size does not change in *dazl* mutant GCs in contrast to wild-type GCs which become smaller.

Interestingly, during cystogenesis the characteristic perinuclear Vasa granules of PGCs (Knaut et al., 2000) are transiently lost and are later reestablished in premeiotic cystocytes. Significantly, zygotic Dazl is not required for Vasa translation, but Dazl protein or successful cyst formation is required to reestablish perinuclear Vasa aggregates. In zebrafish all premeiotic GCs express Vasa, but a subset also express *nanos2;* those that express both are thought to be the GSCs (Beer and Draper, 2013; Cao et al., 2019; Draper, 2017). Based on the observation that all GCs seem to enter a cyst state in wild-type from which Vasa+ premeiotic cells and a limited number of *nanos2+* GSCs emerge, and the finding that GCs of both populations are lost in *dazl* mutants, it is tempting to speculate that the germline cyst not only serves to amplify premeiotic GC numbers but also plays a role in specification of the GSCs. Very recent work in mouse and pig similarly implicates *dazl* in commitment to germline fates (Nicholls et al., 2019), suggesting that whether PGCs, the precursors of the germline are specified by maternal inheritance of germ plasm or are induced later, *dazl* plays and evolutionarily conserved role in commitment or specification of the germline after they reach the gonad anlage.

### Chk2 mediated apoptosis is dispensable for sexual differentiation and fertility

Apoptosis is a feature of zebrafish gonads differentiating as testis (Rodriguez-Mari et al., 2010; Uchida et al., 2002; Wang and Orban, 2007), but the pathways involved in regulating cell death during this process are not understood. Because zebrafish mutants disrupting *tp53* can differentiate as either sex, and mutation of *tp53* suppresses oocyte cell death associated with DNA damage, germline apoptosis is thought to be regulated by both tp53 pathway independent and dependent checkpoints (Bolcun-Filas et al., 2014; Rodriguez-Mari et al., 2010). Here we tested the possibility that the meiotic checkpoint kinase *chk2,* which, in response to DNA damage or aneuploidy, activates both *tp53* and *p63* could account for the *tp53-*independent activity (Abdu et al., 2002; Bolcun-Filas et al., 2014; Sperka et al., 2012). Our observation that mutation of zebrafish *chk2,* as previously observed for *Drosophila* and mouse *chk2* (Abdu et al., 2002; Bolcun-Filas et al., 2014) and zebrafish *tp53* (Bolcun-Filas et al., 2014; Rodriguez-Mari et al., 2010), does not alter viability, sex-specific differentiation, or fertility indicates that normal gonad development occurs independent of Chk2-mediated pathways. Therefore, the pathway that regulates cell death associated with testis differentiation in zebrafish remains to be determined. Based on its ability to suppress death of *dazl* mutant germ cells in mouse mutants, Bax emerges as a compelling candidate (Nicholls et al., 2019).

### Loss of the dazl mutant germline occurs independent of chk2 and p53

Mutation of *chek2* does not disrupt meiosis or fertility in *Drosophila* (Abdu et al., 2002) or mouse (Bolcun-Filas et al., 2014). Similarly, we determined that mutation of zebrafish *chek2* does not interfere with meiosis or fertility in zebrafish. Because *p53* and *chk2,* though not essential for normal development can suppress GC loss in other contexts, we expected that simultaneous mutation of *chk2* or *p53* mutation might similarly suppress germline loss in *dazl* mutants much as *tp53* mutation suppresses oocyte loss and sex reversal of *brca*, *zar1*, *fancl* mutants (Miao et al., 2017; Rodriguez-Mari et al., 2010; Shive et al., 2010). In contrast, although loss of *chk2,* but not *p53,* could prolong survival of abnormal *dazl* mutant GCs, these cells did not progress further in meiosis and eventually were not maintained, resulting in sterility. This is similar to failure of *tp53* mutation to suppress GC loss and sterility of zebrafish *vasa* mutants (Hartung et al., 2014). Like *dazl* mutant GCs, *vasa,* a Dazl target (Haston et al., 2009b; Kee et al., 2009; Li et al., 2019; Reynolds et al., 2005) and *zili* mutant GCs become vacuolated (Houwing et al., 2008). In contrast, *brca, zar1,* and *fancl* mutant oocytes do not and reach later meiotic stages (Miao et al., 2017; Rodriguez-Mari et al., 2010; Shive et al., 2010). Failure of *tp53/chk2* to suppress GC loss in *dazl^-/-^* is consistent with *dazl* function in a premeiotic GC program that acts before and independent of these meiotic checkpoint pathway regulators.

## Author Contributions

Experiments were conceived and designed with contributions from all authors at various stages. AC and FLM generated, identified, and conducted initial analysis of *dazl^ae^* mutant alleles. SB and FLM. TB and JB generated, identified, and conducted initial analysis of the *dazl^Δ7^* allele. SB and FLM performed dissections and performed immunohistological analyses for complementation tests and double mutant analyses. SB performed all other experiments and analysis in consultation with FLM. ER and FLM contributed reagents, materials, and analysis tools. All authors contributed to various aspects of data interpretation and discussion/editing of the manuscript. SB and FLM wrote the manuscript.

## Acknowledgements

We thank members of the Marlow lab for helpful discussions, our animal care staff for fish care (Einstein and CCMS at ISMMS) and the Microscopy CoRE at Icahn School of Mount Sinai. We thank Dr. Derek Stemple and the Zebrafish Mutation Project (ZMP) for providing *chk2^sa20350^* mutants and generating this valuable community resource and ZFIN and ZIRC for curating and making this resource available to the community. Work in the Marlow lab is supported by National Institutes of Health, Grant R01-GM089979 to FLM. MC was supported by an REI fellowship. Work in the Raz lab is supported by the German Research Foundation (DFG), Clinical Research Unit 326, Male germ cells.

