## Supplemental Figures and Legends for "Zebrafish *dazl* regulates cystogenesis upstream of the meiotic transition and germline stem cell specification and independent of meiotic checkpoints"

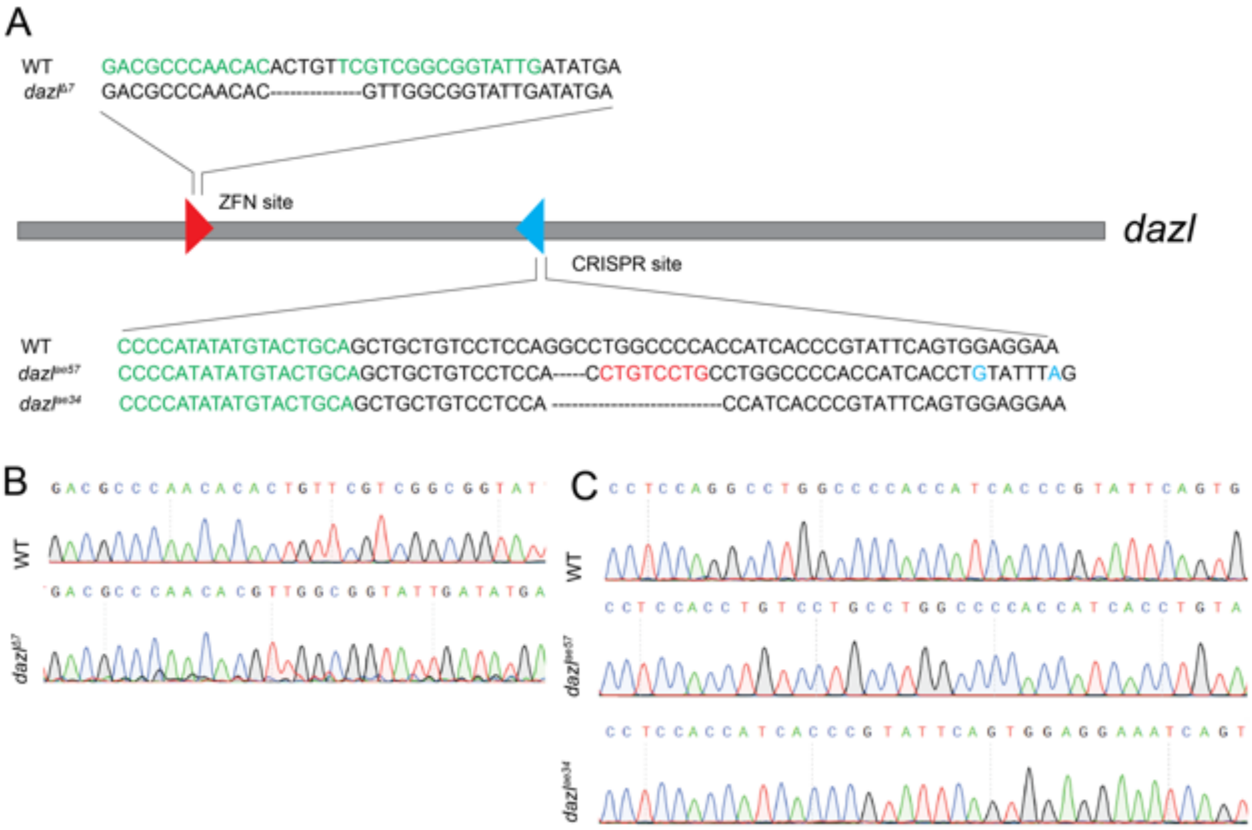


**Supplemental Figure 1. Generation of *dazl* mutants using zinc fingers nucleases and CRISPR-Cas9 mutagenesis.** A Schematic representation of the endogenous *dazl* locus targeted by zinc finger nucleases (ZFN, red triangle) and Crispr-Cas9 (blue triangle) with the target sequences indicated. Both zinc finger nucleases and Cas9 binding sites are highlighted in green. *dazl* alleles generated by ZFN (*dazl*$^{\Delta 7}$) and CRISPR (*dazl*$^{ae57}$and *dazl*$^{ae34}$). Deletions are represented by a dashed line and the substitution is highlighted in blue. (B) Sequence and the corresponding chromatogram of wild type (wt) and *dazl*$^{\Delta 7}$ allele. (C) Sequence and chromatogram alignment of wild-type, *dazl*$^{ae57}$, *dazl*$^{ae34}$ alleles.

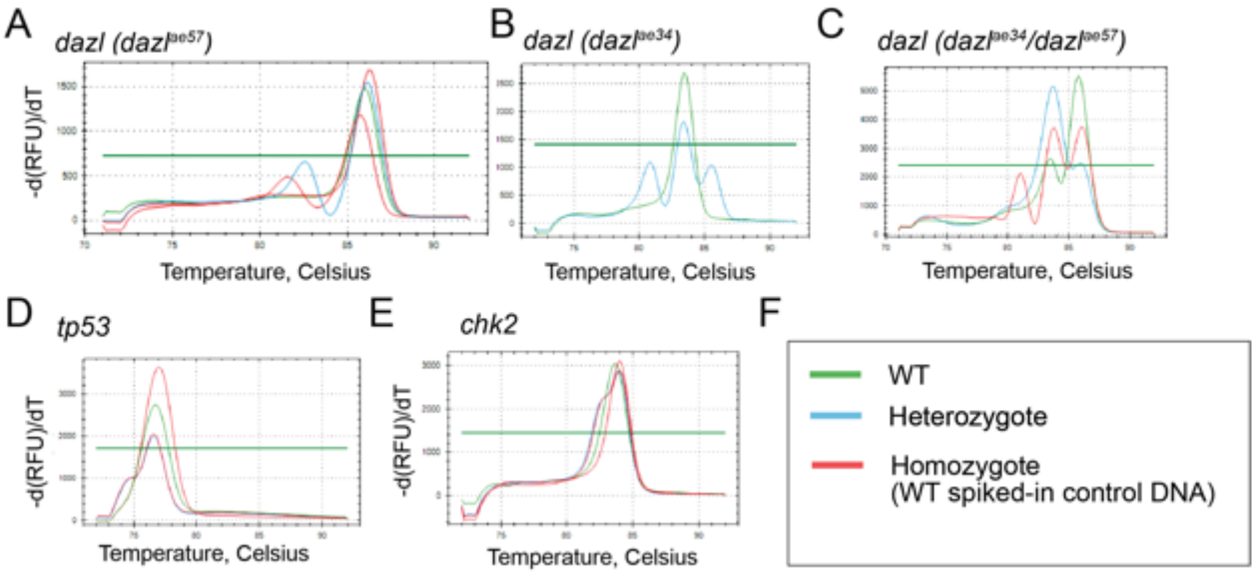


**Supplemental Figure 2. HRMA genotyping assays.** High resolution melt analysis of progeny of heterozygotes intercrosses for $dazl^{ae57}$ (A); $dazl^{ae34}$(B); $dazl^{ae34/57}$(C); *p53* (D) *and chk2* (E). Green, blue lines represent a wild-type or heterozygous genotype respectively. Homozygous mutants are identified by a double red line, which appear heterozygote when wild-type genomic DNA is spiked in a secondary reaction. (F) Legend.

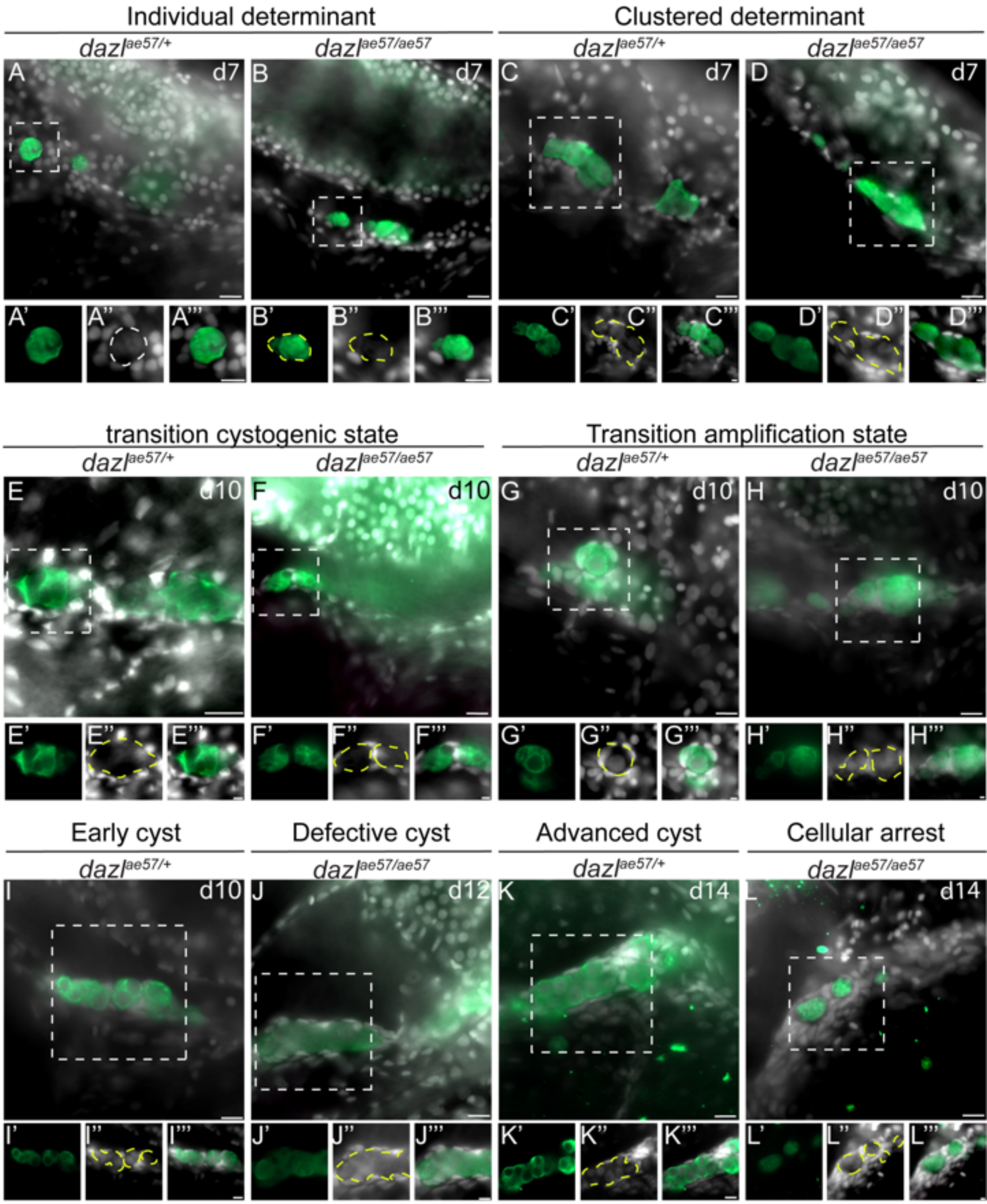

Individual determinant
Clustered determinant
dazl ae57/+
dazl ae57/ae57
d7
transition cystogenic state
Transition amplification state
d10
Early cyst
Defective cyst
Advanced cyst
Cellular arrest
d12
d14

**Supplemental Figure 3. Zygotic *dazl* is required for germline cyst formation and germ cell maintenance.** (A-C''') At 7 and 10d, Vasa labels individual germ cells that are dispersed cells along the germinal epithelium of *dazl*$^{ae57/+}$ or $^{ae57/ae57}$ zebrafish larvae (A-A''' and B-B''', respectively). The boxed cell in A and B are magnified in the bottom inset to show the Vasa signals (green) (A'), DAPI (grayscale), the cell outlines are depicted with yellow dashed lines in the magnified views. Individual PGCs at this stage are characterized by condensed chromatin (white arrowhead), a high nucleus/cytoplasm ratio, and a highly convoluted nucleus surrounded by cytoplasm. Somatic gonadal cells surround the germ cells. Next, individual germ cells group together, termed clustered PGCs hereafter, in wild-type (C-C''') and mutant genotypes (D-D'''). In Clustered GCs Vasa is cytoplasmic but does not appear to be uniform throughout the cytoplasm (C' and D'), and the nucleus becomes more compact (C'' and D''). This step precedes the transition to amplifying divisions and cyst formation (E-E'''-H-H'''). In E and F, synchronous compartmentalized cells, characterized by an enlarged and irregular cell morphology are apparent in wild-type genotypes (E-E''') and in *dazl* mutants (F-F'''). Compartments are apparent as voids in the cytoplasm. During this phase of cyst formation Vasa is diffuse throughout the cytoplasm but granules are not apparent during the transition from individual cells to cystogenesis. Once the 2-cell cyst forms subsequent synchronous cystocyte divisions generate larger cysts in wild-type genotypes (G-G''', I-I''', K-K'''), but not in *dazl* mutants (H-H''', J-J''', L-L'''). The germline cyst cells in wild-type genotypes are characterized by the presence of perinuclear Vasa granules (G'), more uniform chromatin staining (G''). Between 10 to 12d, numerous 2-cell cysts (early cyst) (I) defined by perinuclear Vasa positive aggregates (I'), nuclei symmetrically opposed (I'') fill the gonad. Increasing numbers of somatic gonad cells encapsulate the cysts (I’’’). By 12-14d closely spaced premeiotic cells become apparent (K) among the advanced cysts. Boxed cells are represented in the inset (K’-K’’’). Advanced cysts are defined by the premeiotic germ cell with cytoplasmic and perinuclear Vasa aggregates (K’), and round nuclei with a large nucleolus (K’’), encapsulated by elongated somatic gonad cells (K’’’). In *dazl*$^{ae57/ae57}$ mutants cysts fail to form, instead GCs return to the PGC like morphology and arrest ((H-H’’’, J-J’’’, L-L’’’). bar, 20 μm for the overview and 5 μm for the insets. *dazl*$^{ae57/+}$ (7 d, n=5; 10d, n=7; 12d, n=11; 12d, n=8; 14d, n=6), *dazl*$^{ae57/ae57}$ (7 d, n=4; 10d, n=8; 12d, n=4; 14d, n=4).

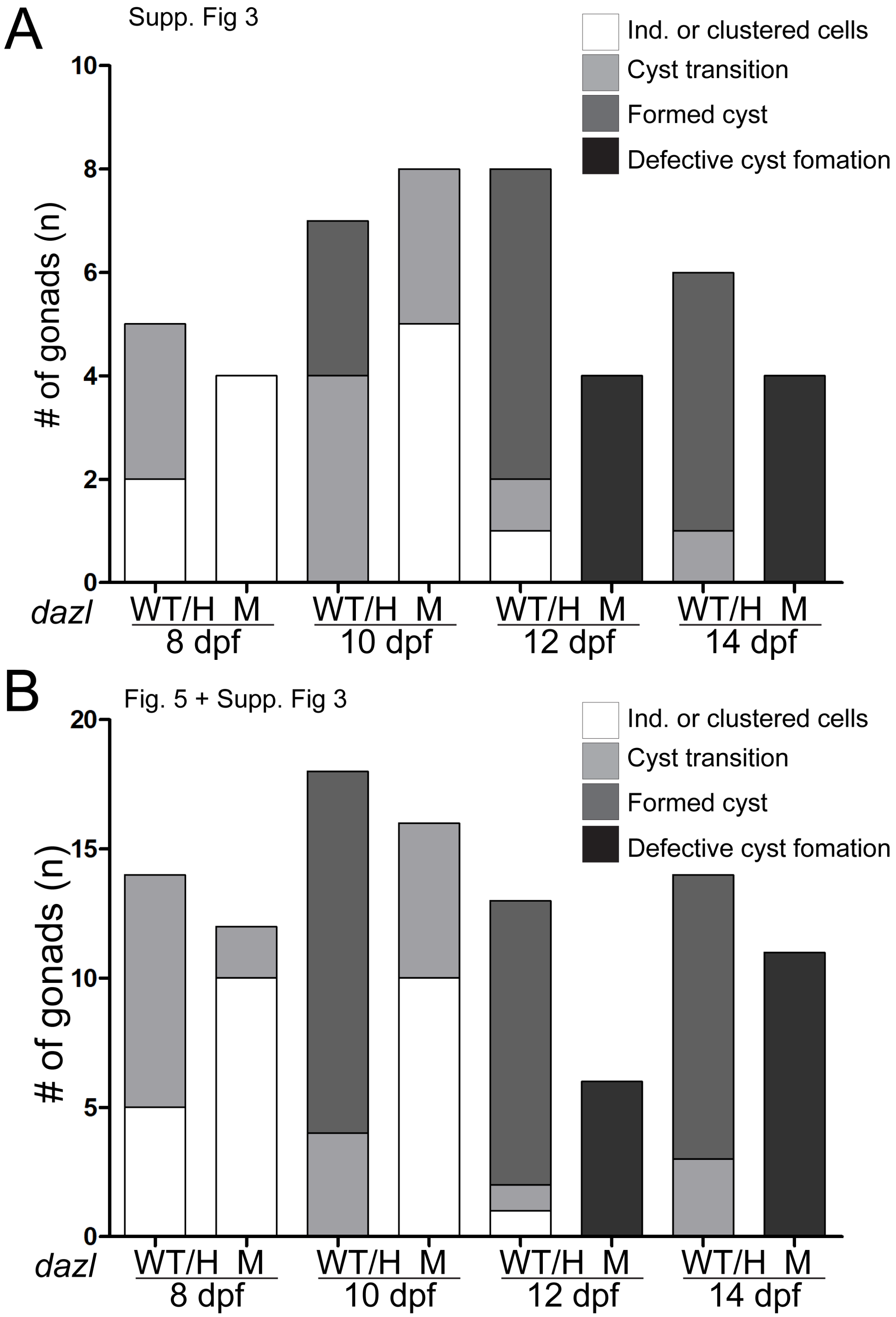
A
Supp. Fig 3
Ind. or clustered cells
Cyst transition
Formed cyst
Defective cyst fomation
### of gonads (n)
10
8
6
4
2
0
dazl
WT/H M
8 dpf
WT/H M
10 dpf
WT/H M
12 dpf
WT/H M
14 dpf
B
Fig. 5 + Supp. Fig 3
Ind. or clustered cells
Cyst transition
Formed cyst
Defective cyst fomation
### of gonads (n)
20
15
10
5
0
dazl
WT/H M
8 dpf
WT/H M
10 dpf
WT/H M
12 dpf
WT/H M
14 dpf

**Supplemental Figure 4. Defective cyst formation in *dazl* mutants.** Bar plots quantifying the gonads analyzed at 8d, 10d, 12d and 14d in wild-type (WT), *dazl*$^{ae57/+}$ and *dazl*$^{ae57/ae57}$ mutants represented in (A) Supplemental Figure 3 and (B) in Figure 5 and Supplemental Figure 3.

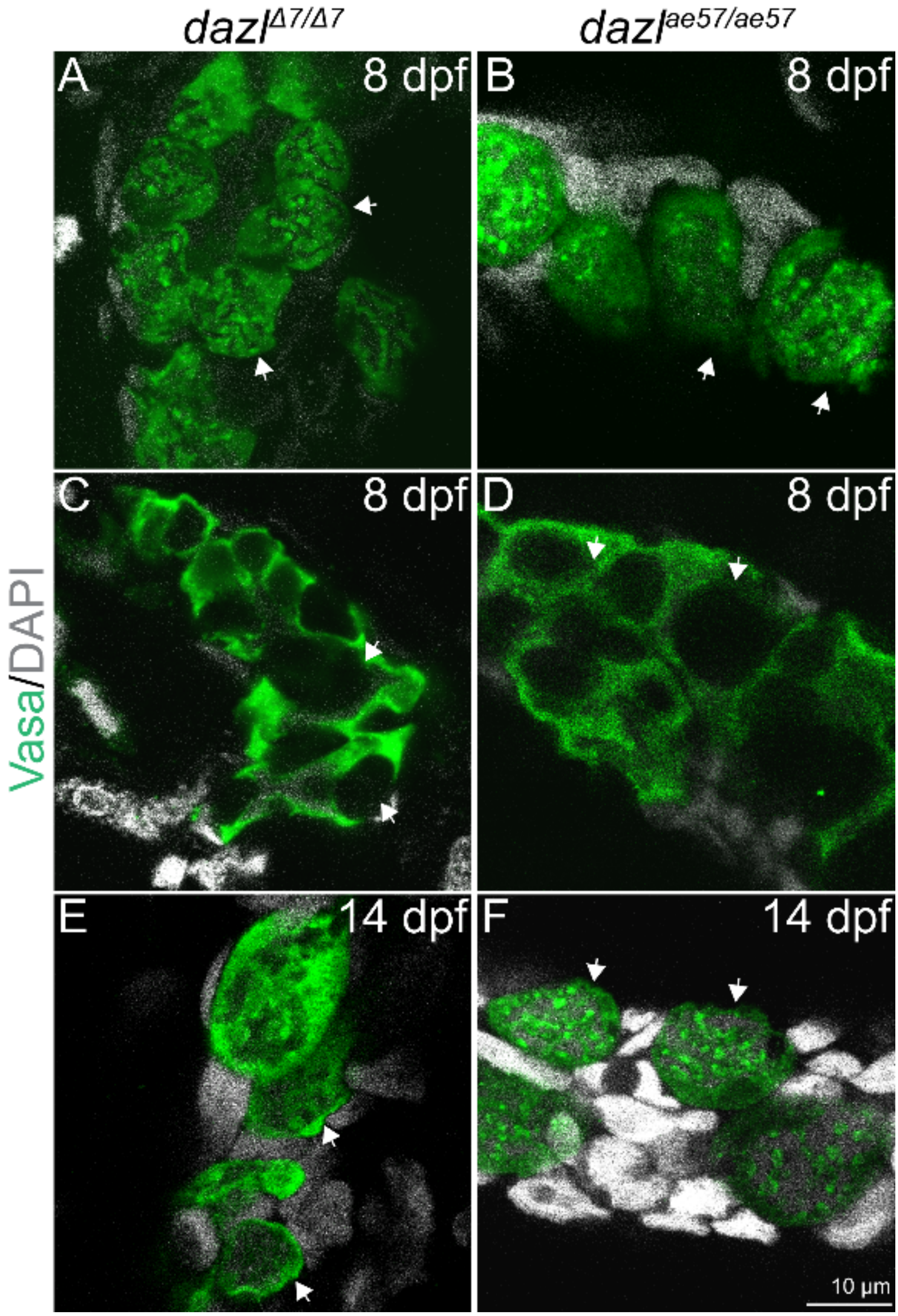
dazl^Δ7/Δ7
dazl^ae57/ae57
A
8 dpf
B
8 dpf
C
8 dpf
D
8 dpf
E
14 dpf
F
14 dpf
Vasa/DAPI
10 µm

**Supplemental Figure 5. Comparison of cystogenesis in *dazl* mutant alleles.** Gonads from *dazl*$^{D7/D7}$ $^{or}$ *dazl*$^{ae57/ae57}$ between 8 and 14d immunostained with Vasa and DAPI. Individual germ cells of (A) *dazl*$^{D7/D7}$ (n=2) and (B) *dazl*$^{ae57/ae57}$ (n=6) mutant gonads. (C, D) Germ cells mutant for either allele initiate cystogenesis as evident from cellular and nuclear morphology, (C) *dazl*$^{D7/D7}$ (n=1) and (D) *dazl*$^{ae57/ae57}$ (n=2). At 14d dazl$^{D7/D7}$ (n=2) (E) or *dazl*$^{ae57/ae57}$ (n=7) (F) mutant germ cells remain as individual cells, indicative of failed cystogenesis.

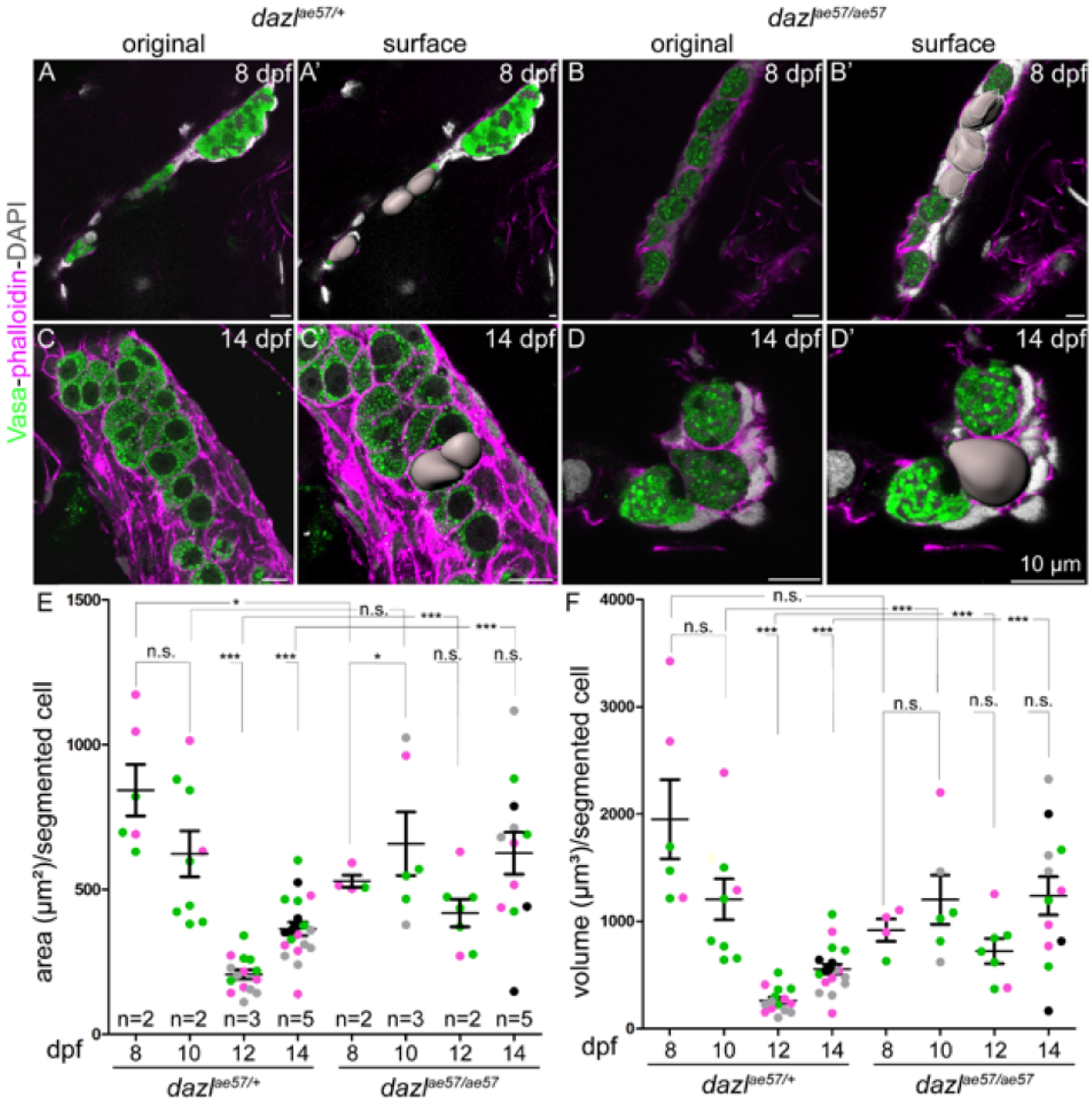


**Supplemental Figure 6. *dazl* mutant cell size between 8 and 14d.** Gonads of *dazl*$^{ae57/+}$ or *dazl*$^{ae57/ae57}$ at 8, 10, 12 and 14d were immunostained with Vasa for GCs; F-actin was stained with phalloidin and nuclei with DAPI. Cells within the Z-stack were manually segmented using Imaris software to measure cell area and cell volume. (A-A') Gonad from *dazl*$^{ae57/+}$ at 8d (A) and the segmented cells in grey. (A-D') Gonads from *dazl*$^{ae57/+}$ at 8d (A) and 14d (C) or *dazl*$^{ae57/ae57}$ (B) and (D). and the respective segmented cells in grey

(A', B', C', D'). (E) Quantification of cell area at 8, 10, 12 and 14d. The number of gonads is indicated at the bottom of each sample points. Each sample value is indicated by the colored dots. (F) Quantification of cell volume at 8, 10, 12 and 14d. The number of gonads is as in (E). Each sample value is differently colored. Two-tailed Student's *t*-test; $*P \leq 0.05$, $***P \leq 0.001$, n.s. not significant. Mean ± s.d.

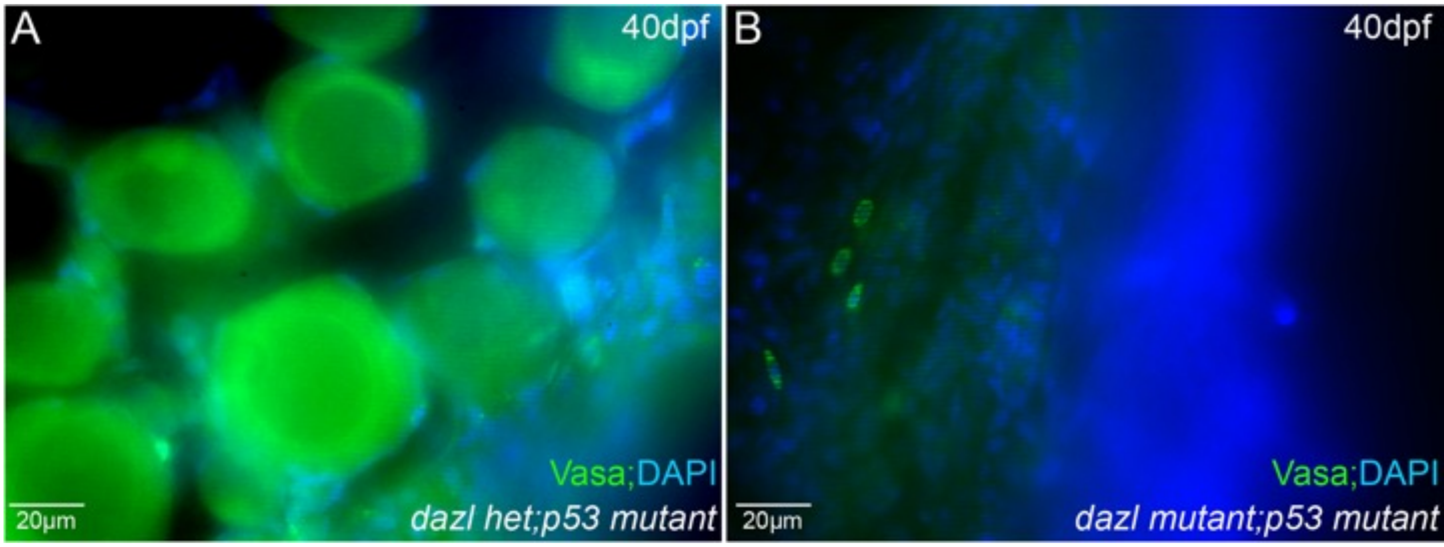


**Supplemental Figure 7. Mutation of *p53* cannot suppress germline loss in *dazl* mutants.** Vasa staining of $dazl^{ae57/+}$ and $dazl^{ae57/ae57}$ in a *p53* mutant background at 40d. (A) Presence of Vasa positive (green) germ cells in a $dazl^{ae57/+}$;*Tp53* mutant gonad. (B) Absence of Vasa positive germ cell in a representative $dazl^{ae57/}$*ae57;Tp53* double mutant gonad.

Bertho et al Supplemental Figure 8

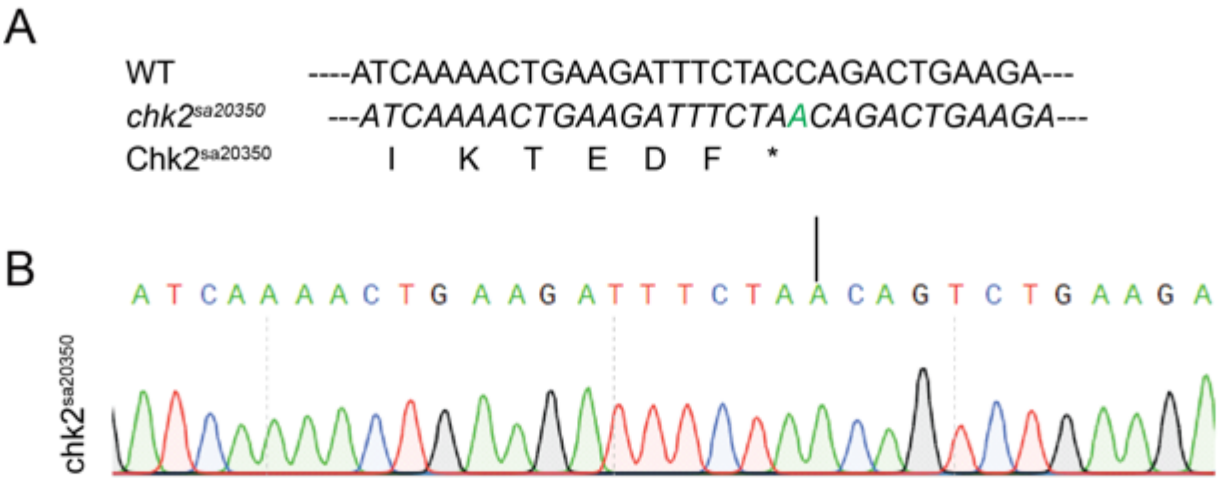


**Supplemental Figure 8. *chk2$^{sa30250}$* is a nonsense allele.** (A) *chk2$^{sa20350}$* sequence at the mutant locus compared to the wild-type (*wt*) and the deduced protein sequence of Chk2$^{sa20350}$. C to A substitution triggers a stop codon. (B) Chromatogram from the sequenced *chk2$^{sa20350}$* locus.
